# Enhancing Perceptual, Attentional, and Working Memory Demands through Variable Practice Schedules: Insights from High-Density EEG Multi-Scale Analyses

**DOI:** 10.1101/2024.07.04.602126

**Authors:** Alexandre Cretton, Kate Schipper, Mahmoud Hassan, Paolo Ruggeri, Jérôme Barral

## Abstract

Contextual interference (CI) enhances learning by practicing motor tasks in a random order rather than a blocked order. One hypothesis suggests that the benefits arise from enhanced early perceptual/attentional processes, while another posits that better learning is due to highly activated mnemonic processes. We propose to harness high-density electroencephalography in a multi-scale analysis approach, including topographic analyses, source estimations, and functional connectivity, to examine the intertwined dynamics of attentional and mnemonic processes within short time windows. We recorded scalp activity from 35 participants as they performed an aiming task at three different distances, under both random and blocked conditions using a crossover design. Our results showed that topographies associated with processes related to perception/attention (N1, P3a) and working memory (P3b) were more pronounced in the random condition. Source estimation analyses supported these findings, revealing greater involvement of the perceptual ventral pathway and the anterior cingulate and parietal cortices, along with increased functional connectivity in ventral alpha and frontoparietal theta band networks during random practice. Our results suggest that CI is driven, in the random compared to the blocked condition, by enhanced specific processes such as perceptual, attentional, and mnemonic, as well as large-scale general processes.

## Introduction

Contextual interference (CI) is a well-known laboratory method for enhancing motor skill learning by implementing variations of a skill within training schedules (Battig 1979; Shea and Morgan 1979; Magill and Hall 1990). A blocked schedule, inducing low CI, necessitates repeated practice of a single skill variation within a block before training other variations in separate blocks. For example, in aiming tasks, a blocked schedule would focus exclusively on one of three variations—such as aiming at one of three different distances—within each block. Conversely, in a random schedule, which induces high CI, the variations of skills are randomized within each block of trials, meaning that aiming at the three different distances would be performed in a mixed and unpredictable order. Laboratory research has extensively reported that random practice typically results in lower performance during training but leads to superior performance in retention and transfer tests (Battig 1978; Shea and Morgan 1979; Magill and Hall 1990).

The benefits of CI on learning are primarily attributed to two sets of hypotheses: the mnemonic (Shea and Zimny 1983; Lee et al. 1985) and the perceptual/attentional hypotheses (Li and Wright 2000; Wright et al. 2016; Lage et al. 2021). The mnemonic hypotheses suggest that the CI effect occurs because increased reliance on working memory processes facilitates long-term memory encoding. For instance, in an aiming task, the random condition may require maintaining all practiced distances in working memory for inter-item comparison or suppressing the action plan of the last practiced distance to facilitate the retrieval of relevant information for the next distance to be executed. On the other hand, the perceptual/attentional hypothesis posits that the dynamic nature of random practice demands increased attention and perceptual processing, which are crucial for adapting to the changing demands of the skills being practiced (e.g. varying distances).

While existing neuroimaging studies support either hypothesis, the heterogeneity in findings—with some studies supporting the mnemonic hypothesis and others supporting the perceptual/attentional hypothesis—may stem from the absence of studies combining sufficient spatial and temporal resolution to investigate the dynamic processes occurring within short time windows. In the remainder of this introduction, we will examine neuroimaging studies supporting each hypothesis. Subsequently, we propose an experiment utilizing the spatio-temporal capabilities of high-density EEG in a multi-scale analysis approach to better understand the dynamics of processes involved in CI. Our goal is to gather evidence that may help reconcile the mnemonic and perceptual hypotheses regarding CI.

Neuroimaging studies provide indirect evidence supporting the mnemonic hypotheses by demonstrating greater engagement of executive function regions (e.g., prefrontal cortex (PFC)) during random practice than during blocked practice (Lage et al., 2015; Wright et al., 2016). PFC is known to be involved in maintaining and manipulating information in working memory (Constantinidis and Klingberg 2016; Riley and Constantinidis 2016), selecting relevant information, and executing functions essential for cognitive processing (Lara and Wallis 2015; Friedman and Robbins 2022). Kantak et al. (2010; 2011) demonstrated that applying repetitive Transcranial Magnetic Stimulation (rTMS) over the Dorsolateral Prefrontal Cortex (DLPFC) during skill acquisition impaired one-day skill retention in the random group but not in the blocked group. In a series of fMRI experiments conducted by Lin and colleagues (Lin et al. 2011; Lin et al. 2013; Lin et al. 2016), participants performed two training sessions on a Serial Reaction Time (SRT) task under either random or blocked schedules. They were then retested three days later on the same task. Lin et al. (2011) observed that, during the acquisition phase, the random group showed higher activity in the occipital/temporal cortex, sensorimotor regions, and PFC compared to the blocked group. However, during the retention phase, this activity pattern was reversed; the random group exhibited lower activation in the medial prefrontal, premotor, and parietal areas. Furthermore, Lin et al. (2013) reported enhanced functional frontoparietal connectivity between the DLPFC and superior medial frontal regions, supplementary motor area, caudate nucleus, and both the inferior and superior parietal lobes in the random condition during the second training session. This increased connectivity, which persisted during retention, correlated with the superior retention benefits observed following random practice.

The alternative hypothesis that focuses on attentional and perceptual mechanisms, is also supported by neuroimaging data. For instance, Lelis-Torres et al. (2017), while exploring the dynamics of perceptual and working memory processes in a sequence pressing timing task, observed that an EEG marker of task engagement, encompassing perceptual and attentional processes, decreased during random practice compared to blocked practice. Conversely, a second EEG marker of mental workload, which involves working memory processes, exhibited no significant difference between the random and blocked practice conditions. Additionally, in an experiment in which participants engaged in a bimanual visuomotor tracking task during an fMRI scan, Pauwels et al. (2018) observed that random practice, compared to blocked practice, recruits regions important for visual processing, such as the right MT/V5 and bilateral LOC. Moreover, in the same protocol, Chalavi et al. (2018) reported a decrease in GABA+ levels in the occipital cortex following random practice. This decrease suggests neurochemical changes in the excitatory-inhibitory balance, favoring the decreased inhibition of perceptual processing mechanisms under random conditions. In a study that inferred perceptual and attentional effort through eye tracking in a sequence-pressing timing task, participants in the random practice condition exhibited significantly higher levels of pupil dilation and increased frequency of eyeblinks during the acquisition phase (Bicalho et al. 2019). The authors interpret these physiological indicators as suggestive of increased perceptual effort required during the initial learning phase in random practice. Finally, Frömer and colleagues (2016) trained participants in a virtual dart game, where the target could appear centrally, to the left, or to the right of the screen. The study observed a tendency towards a greater decrease in P3 amplitude, an ERPs associated with attentional resources allocation, within a 250-400 ms window at prefrontal, frontal, and central regions of interest (ROI) during random practice compared to blocked practice.

Both mnemonic and perceptual/attentional hypotheses have been supported by several studies using various neuroimaging methods, each contributing to our understanding of CI. However, the different strengths and limitations of these methods have influenced the ability to support both hypotheses within the same studies. For instance, fMRI is renowned for its excellent spatial resolution but is limited by its low temporal resolution due to its reliance on slow blood-oxygen-level-dependent (BOLD) signals ( Logothetis et al. 2001; Heeger and Ress 2002). This limitation poses challenges in CI studies that involve multiple-stage processes (Immink and Wright 1998; Immink and Wright 2001; Wright et al. 2004), as distinguishing between stages such as response selection and movement programming (Mulder et al. 1984; Klapp 1995; Spijkers 1990; Masaki et al. 2004), which occur within tens of milliseconds (Mulder et al. 1984; Masaki et al. 2004). Conversely, EEG excels in temporal resolution by directly recording real-time electrical fluctuations, allowing for precise monitoring of neural dynamics on a millisecond scale (Jorge et al. 2014; Turner et al. 2016). This makes EEG particularly effective for investigating rapid event-related potentials (ERPs) related to attention, perception, and working memory. For example, the N1 component indicates heightened visuospatial attention (Krigolson et al. 2015), the P3a component is associated with frontal attention mechanisms (Polich 2007), and the P3b component relates to working memory processes (Polich 2007). Despite the advantages of EEG, studies supporting the perceptual hypothesis have not fully utilized its temporal resolution, often relying on spectral analyses (Thürer et al. 2017; Beik et al. 2020; Beik et al. 2022), which are less effective for investigating rapid neural dynamics. Additionally, these studies frequently limit their analyses to predefined regions of interest (ROIs) and specific timings, thus not fully integrating EEG’s temporal and spatial capabilities (Frömer et al. 2016; Lelis-Torres et al. 2017).

To advance our understanding of CI, a multi-scale approach combining the spatial precision of fMRI with the temporal accuracy of EEG could be highly beneficial. This integrated approach would allow for a more comprehensive examination of the dynamic processes underlying attention and memory, providing insights that single-method studies may not achieve. To achieve this goal, we developed a computer-based aiming task requiring participants to make straightforward movements to click on a target from three different starting positions while recording high-density EEG. This task, structured as a crossover design, had 35 participants complete the distances in both blocked and random orders without any visual feedback of the effector’s position or the mouse cursor. During the movement preparation period, we used scalp topographic analyses to identify periods where neural generators differed between conditions. We then performed source estimation analysis to localize the specific cortical neural generators responsible for these topographical differences. Finally, we computed large-scale functional connectivity to explore network differences between conditions.

Drawing on research supporting mnemonic or perceptual/attentional hypotheses, we hypothesize that brain-electrocortical markers indicative of working memory engagement and attentional demand will be more pronounced in the random condition compared to the blocked condition. Specifically, we expect this to manifest at the N1 and P3a components for attentional processes, and at the P3b component for working memory processes. Additionally, we anticipate that source analysis will confirm greater activity in attentional- and mnemonic-related areas. Finally, in the random compared to the blocked condition, we expect the functional connectivity analysis to show significant interconnections within the frontoparietal Executive Control Network (ECN), which is crucial for coordinating high-level cognitive processes such as attention regulation, problem-solving, and working memory.

## 2 MATERIAL AND METHODS

### 2.1 Participants

We recruited 36 right-handed students from the University of Lausanne (12 males; mean age: 20.9 years, standard deviation: 2.31 years; age range: 18-28 years). One participant was excluded due to an EEG recording affected by unmanageable slow wave artifacts (the participant reported being very tired), resulting in a sample size of 35 participants (11 males; mean age: 20.9 years, standard deviation: 2.34 years; age range: 18-28 years). Participants gave written informed consent, met the eligibility criteria, which included having normal or corrected-to-normal vision, and reported no history of psychiatric or neurological disorders. We provided participants with compensation of 100 Swiss Francs (approximately 107 US dollars at the time) for their participation. The local ethical committee approved all procedures before the beginning of the study (CER-VD: 2021-01456).

### 2.2 Task and material

The task (Fig 1A) required participants to aim at a target with a computer mouse using their left non-dominant arm. By moving the arm forward, the cursor of the mouse was displaced from a starting point at the bottom of the computer screen to a target at the top. Both the target and starting positions were represented by a one centimeter wide and high cross (vertical and horizontal cross dimension in degree of visual angle: 0°49’). Participants had to execute a fast and straight-forward movement and click with the mouse button as close to the target as possible. In each trial, the distance between the starting point and target could either be 7 cm (short distance), 14 cm (mid distance), or 21 cm (long distance). The corresponding vertical visual angles were 5°43’, 11°25’ and 17° 30’, respectively.

**Figure 1.**
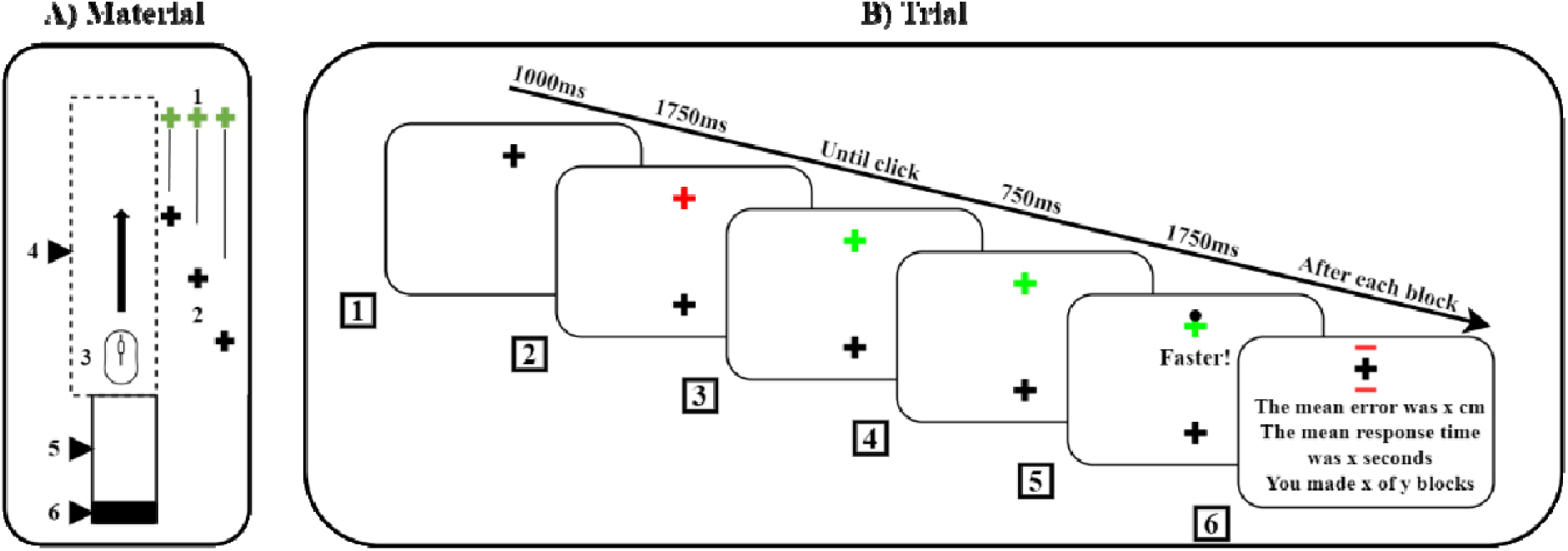
Material and Trial. A) Material: The experimental setup included (1) the target, (2) starting points at short, mid, and long distances, (3) an ambidextrous computer mouse, (4) a wooden box obscuring the hand, (5) a wooden support for the arm, and (6) a wooden wedge brace for the elbow during the inter-trial interval. The computer monitor displaying the task is not depicted in this representation. (B) Trial: (1) A trial began with a fixation cross displayed for 1000ms. (2) Following this, participants prepared to move for 1750 ms when two crosses appeared simultaneously: a black starting point and a red target. (3) Once the target changed color from red to green, (4) participants swiftly moved the invisible cursor from the starting cross (located at the bottom of the screen) to the target cross (at the top) and attempted to click as close to the target as possible. (5) A 750ms pause ensued, after which feedback was shown for 1750 ms—a black dot indicating the cursor’s click position. (6) If the response time surpasses 20% of the baseline established in the familiarization trials, the black dot would not appear; instead, a prompt reading “faster” would be displayed. After each block, participants received summary feedback, which included red outlines around the target—representing the mean error in centimeters— and mean response time (in seconds) along with information about the number of completed and remaining blocks.

At the outset of the task, the participants completed a familiarization phase where they performed six trials of each distance. To standardize movement execution across trials, their forearms rested on a wooden support with their elbows blocked by a wooden wedge. At the beginning of each trial, the forearms formed a 90-degree angle with the upper arm, which was parallel to the body. To minimize head movement during the experiment, participants used a chin rest device. This ensured that their eyes remained fixed in the middle and 70 cm away from the screen, providing constant visual access to both the starting point and target without the need for eye movement. The position of the wooden support and the height of the seat were adjusted for each participant. Additionally, participants could not initiate cursor movement prior to the start of the trial, as the cursor was automatically set and blocked to the starting position to prevent any anticipatory movement. The cursor was constrained to move along the y-axis only and could not deviate along the x-axis. To prevent visual feedback and online control of movement, neither the cursor’s displacement nor the hand displacements were visible during the pointing movements.

The task was powered by the PsychoPy 2021.1.2 (Peirce et al. 2019) software and displayed on a Dell Alienware AW2521HFA computer monitor. The monitor had a refresh rate of 240Hz, a screen resolution of 2560 x 1440 pixels, and a 24.5-inch screen size. The monitor’s IPS response time of 1 ms (grey to grey) enabled smooth and precise cursor movements. Arm movements were executed using an ambidextrous computer mouse (Kova AIMO), which was set to a resolution of 200 dpi to control the cursor.

### 2.3 Trial

The chronological sequence of one trial during the test session is depicted in Figure 1E. At the beginning of each trial, an intertrial fixation cross was displayed on the screen at the target location for 1000 ms. Then, two crosses, one indicating the starting position (in black) and the other one (in red) the target to hit were presented during 1750 ms. Participants were required to move towards the target when the color of the target cross changed from red to green. The movement execution period remained open until the participant clicked the mouse button in the vicinity of the target. After 750 ms, an immediate feedback – a black dot – appeared on the screen at the location of the click. If the total trial time (i.e., reaction time + movement time) exceeded 20% of the average time computed during familiarization (performed at the beginning of each task), the black dot feedback was replaced by a message “faster” at the center of the screen. The black dot or the message were displayed for 1750 ms. Finally, at the end of each block, feedback displaying the mean values of error and response time was presented along with the number of completed and remaining blocks. Simultaneously, a graphical representation of the mean error drawn around the target was displayed (red lines on the last panel of Figure 1E), allowing participants to self-evaluate their performance. At this moment, they could choose to take a break or move to the next block by pressing the space bar.

### 2.4 Experimental protocol

The task lasted 25 minutes. The task comprised 216 trials, organized into 9 blocks of 12 trials per experimental condition (i.e., blocked and random). In the blocked condition, each block exclusively focused on one of the three distances. This resulted in an order of three consecutive blocks for each distance (e.g., three blocks for the mid distance, followed by three blocks for the short distance, and then three blocks for the long distance). In the random condition, each block included 12 trials split into 4 trials for each distance, presented in a random order. It was a crossover design as each partic pant completed each condition. Both the order of the conditions and the order of the distances in the blocked condition were counterbalanced across participants. It’s important to note that the datasets used in this study are extracted from a training protocol conducted over three consecutive weeks, comprising nine training sessions. The blocked and random conditions were tested one day before the training sessions (baseline), the day after last training session, and one week later. The tests included both the trained task and a transfer task, which involved the untrained hand and new distances. For the current study, we analyzed behavioral and EEG collected during the baseline test.

### 2.5 Metrics of performance

#### Errors

Errors were quantified with two metrics: absolute error (AE) and variable error (VE). AE is the average absolute distance of endpoint clicks relative to the target, regardless of the direction of the deviation. VE is the standard deviation of the endpoint clicks, regardless of the target position. Higher accuracy and consistency in aiming movements are indicated by smaller AE and VE values, respectively (Fitts 1954; Schmidt et al. 1979).

#### Speed

To compute speed, we measured the distance covered by the hand (in centimeters) until the click and divided it by the movement time (interval time between the onset of movement and the click, in seconds). Speed is often inversely linked to accuracy (Fitts 1954) and consistency (Schmidt et al. 1979) of movement. We quantified the mean speed (MS), which is the average speed across all trials, and the trial-to-trial speed variability (VS), defined as the standard deviation of the speed across all trials.

#### Reaction time

Reaction time, measured in seconds, is the interval between the moment the cross changes from red to green and the onset of movement, i.e., the first timeframe where the movement of the mouse is detected. It is defined as the time required for an individual to prepare and initiate a movement in response to a stimulus (Munro et al. 2007). We computed the mean reaction time (MRT) across all trials, and the trial-to-trial reaction time variability (VRT), defined as the standard deviation of reaction times across trials.

#### Behavioral data exclusion and statistical analyses

For the behavioral analyses we removed trials that were initiated before the target cross turned green and trials with elapsed response delay (i.e., exceeding 20% of the response time computed in the familiarization trials). Then, we modelled a linear mixed model (LMM) for each of the six behavioral variables (i.e., AE, VE, MS, VS, MRT, VRT) with condition (blocked, random), and distance (short, mid, long) as fixed factors and subjects as a random factor (random intercept). We tested the main and interaction effects on the response variables. In case of a significant interaction (p<0.01), we computed pairwise comparisons and applied Holm-Bonferroni corrections to control for type-I errors. Prior to analysis, we log-transformed all response variables to meet the normal distribution assumption of the statistical models used. The statistics in the results section are based on log-transformed data. However, to make the statistical differences more intuitively understandable, we present the deltas in their non-log-transformed values. The results of the Kolmogorov-Smirnov normality test are presented in Supplementary Material (Table S1). The statistical analyses were performed using Jamovi 2.3.18 (The jamovi project 2023).

### 2.6 EEG

We recorded continuous EEG during the task at a rate of 1024 Hz with 24-bit A/D conversion using 128 electrodes (Biosemi ActiveTwo system) disposed according to the international 10–20 system. Two additional electrodes were used (active CMS: common-mode sense and passive DRL: driven right leg) to form a feedback loop for the amplifier reference. Brain Vision Analyzer (BrainVision Analyzer, Version 2.2.2, Brain Products GmbH, Gilching, Germany) was used for the preprocessing steps. The raw EEG signals were filtered between 0.5 and 45 Hz with 50 Hz notch using a zero-phase shift second-order Butterworth filter, and downsampled to 512 Hz. Eye blinks and saccades artifacts were corrected using independent component analysis (ICA) (Cardoso 1998) and bad electrodes replaced using linear spline interpolation of adjacent electrodes (Perrin et al. 1987). The dataset was segmented into epochs of 2500 ms (200 ms before warning cross-stimulus to 550 ms after imperative cross-stimulus). We excluded epochs containing electrical potential values exceeding ±80 μV, in addition to trials removed at the behavioral level. We then recalculated the data to obtain an average reference. After preprocessing, the mean number of remaining trials was 85.86 (79.52%) +/- 13.34 for the random condition and 85.91 (79.55%) +/- 10.56 for the blocked condition. No baseline correction was applied.

#### 2.6.1 Consistency test, topographical analysis of variance and GFP analysis

We computed an event-related potential (ERP) for each condition in the BrainVision software by averaging the epochs from each subject. Subsequently, we conducted ERP analyses using RAGU (Randomization Graphical User interfaces) software implemented in MATLAB (http://www.mathworks.com/) (Koenig et al. 2011; Habermann et al. 2018). To evaluate the results, we performed non-parametric randomization statistical tests, including the topographic consistency test (TCT), topographic analysis of variance (TANOVA), and global field power (GFP) analysis (Koenig et al. 2011; Habermann et al. 2018). All analyses were conducted on the 2500 ms epochs using 5,000 randomization runs, with a p-value threshold set at 0.01 to ensure robustness against Type I errors. To further safeguard against false positives due to multiple comparisons, for TANOVA and GFP, we employed additional testing based on the duration of significant time periods exceeding chance. This procedure enabled us to confirm that the observed significant effects were not random variance artifacts. Similar procedures employing RAGU have proven effective in identifying neural electrocortical correlates in differents contexts (Ruggeri et al. 2019; Ruggeri et al. 2020; Di Muccio et al. 2022; Simonet et al. 2022; Raynal et al. 2024).

We conducted the TCT separately on each of the two ERPs to identify time intervals characterized by a consistent pattern of active sources (i.e., topographically similar scalp potential distribution) among participants. This ensured that the stimulus presentation consistently activated common neuronal sources across subjects over time, thereby minimizing the risk of drawing incorrect conclusions from TANOVA and GFP analyses (Habermann et al. 2018).

TANOVA analysis is a non-parametric randomization test that evaluates global dissimilarities in electric field topographies without requiring the pre-selection of specific time points for event-related potential (ERP) analysis. Unlike channel-wise comparisons that identify differences at individual electrode levels, TANOVA assesses global dissimilarity across the entire electric field topography between conditions, testing for significant topographic distinctions at each time point. This methodology enables the identification of significant differences resulting from variations in the active sources of the evoked potentials, rather than mere differences in source strength. To this end, the analysis was performed on amplitude-normalized maps (Global Field Power, GFP = 1) to ensure that observed topographical differences were not influenced by global field strength.

GFP analysis quantitatively assesses the overall strength of neural activation between the “blocked” and “random” conditions. GFP is the standard deviation of all EEG electrode potentials at a single time point, providing a single value that represents the total electric field strength at that moment. GFP analysis quantifies overall electric field power without considering its spatial distribution. This is distinct from TANOVA, which investigates the spatial patterns of neural activation by examining the dissimilarity in electric field topographies across the scalp. Together, GFP and TANOVA provide a holistic view of neural dynamics, with GFP contributing to a measure of neural activation strength and TANOVA detailing the spatial arrangement of this activity.

Finally, we classified the dynamic changes in ERP topographies through microstate analysis, which decomposes the ERP signal into a set of stable topographical map configurations called microstates. For this analysis, we assumed that the topographies might vary between groups in their duration. We computed the microstate clustering using the algorithm implemented in RAGU. First, the algorithm identifies the optimal number of microstate prototype maps through a cross-validation procedure—an iterative technique that requires randomly splitting the available ERPs into learning and testing sets to determine their topographical patterns. We used the k-means algorithm with 100 random initializations to identify each microstate map. Final microstate maps were computed using the grand average ERPs during the blocked and random condition in the interval from 156 to 687 ms where we identified significant condition differences with the TANOVA and GFP analyses (described in the result section).

### 2.6.2 Source analysis

For the periods marked by significant differences we employed the weighted minimum-norm estimation (WMNE) method (Baillet et al. 2001) for neural source estimation, utilizing Brainstorm software (Tadel et al. 2011). This approach offers a stable and conservative inverse solution that is widely applicable across various studies (Baillet et al. 2001). Given the absence of individual MRI templates, we constructed a generic head model using the OpenMEEG Boundary Element Method (BEM) (Kybic et al. 2005; Gramfort et al. 2010). This model assumes standard conductivities for the scalp and brain layers (1.0000 S/m) and assigns a conductivity of 0.0125 S/m for the skull layer. We applied the WMNE method assuming independent noise across electrodes (noise covariance modeled with an identity matrix) to facilitate a stable inverse solution, alongside fixed source orientation and depth weighting (exponent of 0.5 and a weight limit of 10) to accurately localize neural activity. We set the regularization parameter at a noise level of 0.1 (Baillet et al. 2001). For the source comparison between conditions in each of the significant periods, we conducted statistical comparisons using independent parametric tests, with a significance level set at p<0.01 and adjustments for multiple comparisons through the False Discovery Rate (FDR) method (Tadel et al. 2019).

### 2.6.3 Connectivity Analysis

To analyze the connectivity differences between the random and blocked conditions, we initially calculated phase-locking value (PLV) matrices utilizing Brainstorm software (Tadel et al. 2011). This involved transforming EEG data into source space using the Desikan-Killiany atlas (Desikan et al. 2006), made of 68 cortical regions. Such delineation facilitates a detailed analysis of connectivity patterns across the cortex by evaluating the synchronization among these regions, represented in a 68×68 connectivity matrix. PLVs were computed within specific frequency bands after decomposing the EEG signals through the Hilbert transform. This approach enabled us to assess phase synchronization in the theta (4-7 Hz), alpha (8-12 Hz), beta (13-30 Hz), and gamma (30-45 Hz) frequency bands. We introduced a conservatively applied window of at least three cycles of the lowest frequency analyzed around each period identified as significant by TANOVA and GFP analyses: 750 ms for theta, 375 ms for alpha, 230 ms for beta, and 100 ms for gamma around each window. However, to ensure the interpretability of the data, we adjusted the windows to avoid overlaps and to prevent them from being split in the middle. PLVs were calculated for each trial within the selected time windows and for each of the selected frequency bands. We then concatenated these PLVs to derive a single PLV value per subject and condition for each frequency band. Consequently, for each considered time window and frequency band, every subject received a PLV for both the random and blocked conditions.

To examine connectivity variations between random and blocked conditions, we conducted directional paired t-tests using Network-Based Statistic (NBS) software (NBS toolbox, https://www.nitrc.org/projects/nbs/) (Zalesky et al. 2010) on PLVs across selected time windows and frequency bands. We conducted the analysis for both directions: random > blocked and blocked > random. A significance level of p < 0.01 was set, employing 5000 permutations, and controlling for multiple comparison with the family-wise error rate (FWER) reducing he probability of false networks identification.

## 3. RESULTS

### 3.1 Behavioral results

Figure 2A shows that AE scores were significantly higher in the random condition (M = 2.300, SD = 0.841) compared to the blocked condition (M = 1.770, SD = 0.769), F(1, 172) = 82.400, p < .001. Similarly, Figure 2B indicates that VE scores were significantly influenced by condition, with higher scores in the random condition (M = 1.680, SD = 0.570) than the blocked condition (M = 1.370, SD = 0.661), F(1, 172) = 55.200, p < .001). In contrast, Figure 2C illustrates that MRT scores did not show a significant difference between conditions (Blocked: M = 0.274, SD = 0.064; Random: M = 0.267, SD = 0.060), F(1, 172) = 3.140, p = 0.780. VRT scores, shown in Figure 2D, were not significantly different between conditions (Blocked: M = 0.097, SD = 0.036; Random: M = 0.098, SD = 0.037), F(1, 174) = 0.150, p = 0.699). However, MS scores presented in Figure 2E were significantly higher in the blocked condition (M = 33.600, SD = 17.700) compared to the random condition (M = 32.200, SD = 18.300), F(1, 172) = 17.200, p < .001. Finally, Figure 2F shows that VS scores were significantly higher in the random condition (M = 7.460, SD = 3.750) compared to the blocked condition (M = 6.910, SD = 3.580), F(1, 172) = 8.920, p = 0.003).

**Figure 2.**
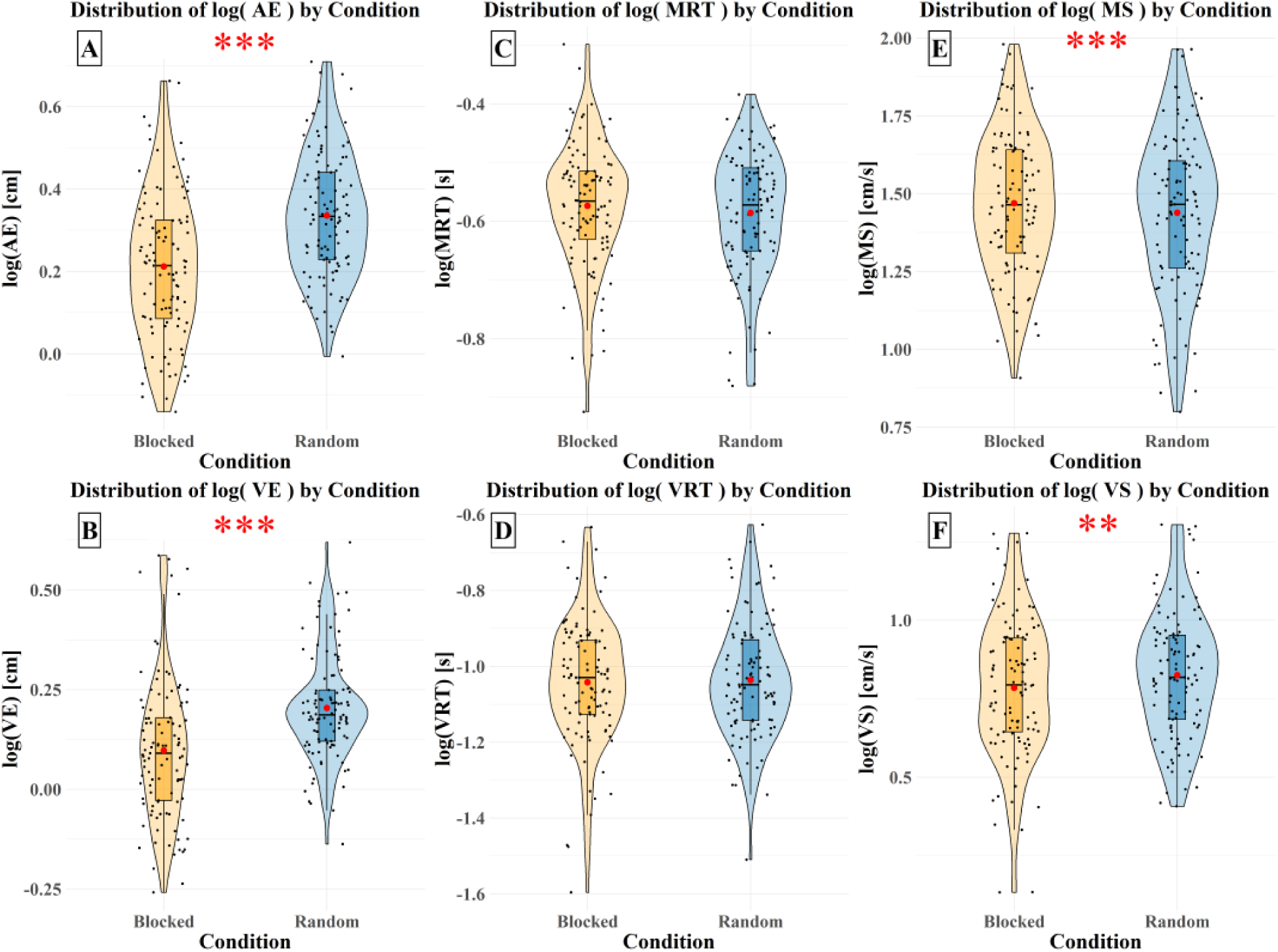
Logarithmically transformed data distribution of AE (A), VE (B), MRT (C), VRT (D), MS (E), and MS (F) across blocked (orange) and random (blue) conditions. The red points represent mean values. Significance: **(p ≤ 0.01), ***(p ≤ 0.001).

### 3.2 Topographical analyses

The TCT, illustrated in the Supplementary Material (Figure S1), was applied to event-related potentials (ERPs) from both blocked and random conditions. This analysis revealed periods of consistent topography across subjects. In the blocked condition, consistent topographies were observed between - 110 ms and 812 ms and from 1181 ms until the end of the epoch. In the random condition, consistent topographies were maintained from -69 ms to 1093 ms, and from 1522 ms to the end of the epoch. According to the consistent EEG periods identified by the TCT, significant differences in topography between these conditions were detected using TANOVA within the time periods 281–404 ms and 434–552 ms (Figure 3A, time periods highlighted in yellow), and significant power differences were identified using GFP analysis within the time periods 156–222 ms, 457–542 ms, and 559–687 ms, all confirmed by the global duration post hoc test (Figure 3B, time periods highlighted in purple).

**Figure 3.**
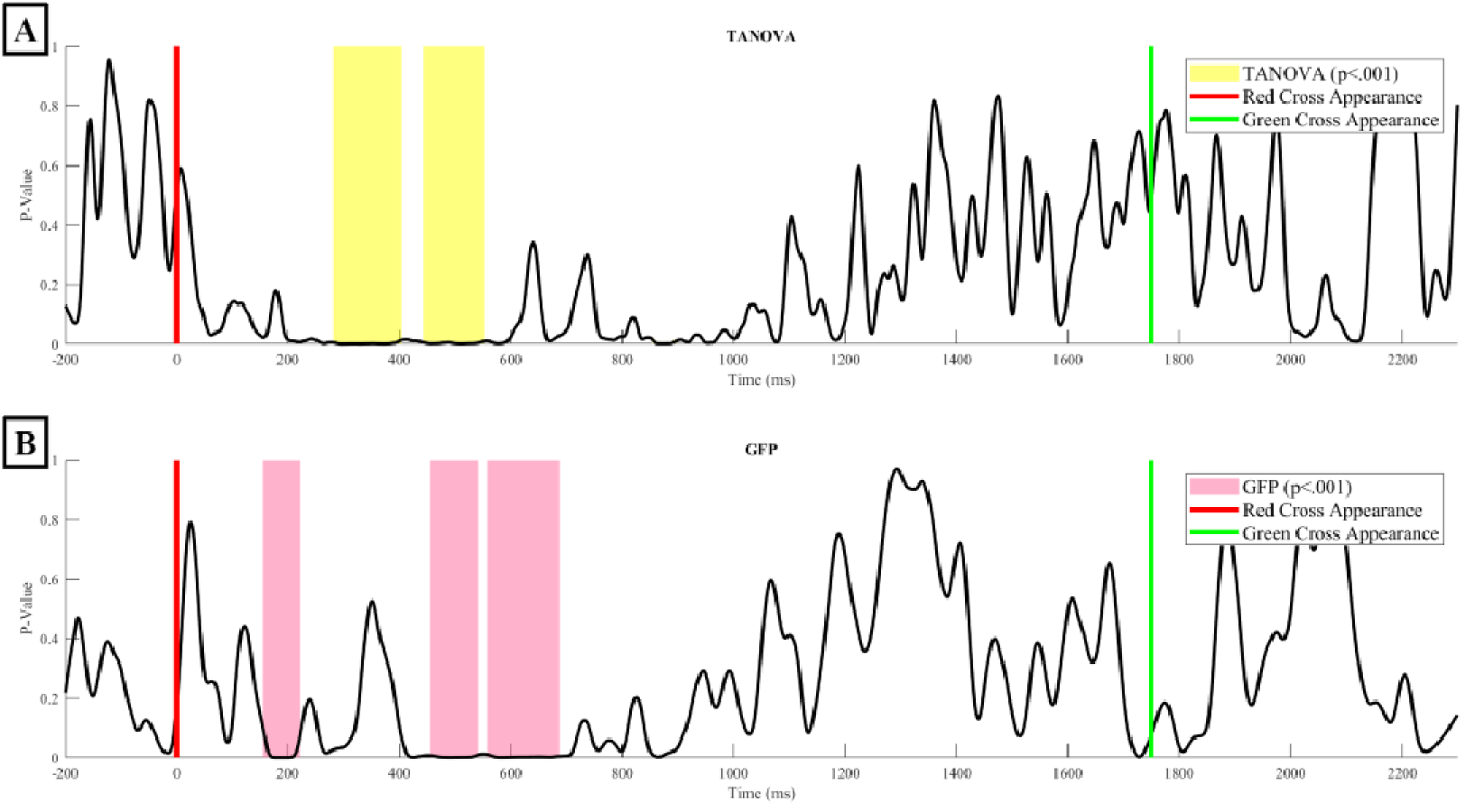
Results of TANOVA (A) and GFP (B) analyses, displaying the p-values of the differences between conditions over time. Yellow shaded areas indicate periods where TANOVA’s p-value falls below p<.001, which survive post-hoc tests on duration. Pink shaded areas indicate periods where GFP’s p-value falls below p<.001, which survive post-hoc tests on duration. The red line indicates the moment when the distance is displayed, and the green line represents the imper tive stimulus signaling participants to aim at the target.

For subsequent analyses, we focused on the 156–687 ms interval, which presented consistent topographies across subjects and showed periods of significant topographical differences. First, we conducted microstate analyses to identify patterns of brain activity and temporal dynamics across conditions (Figure 4). The sequence of maps is consistent across both conditions for the first two maps but diverges thereafter (Figure 4). Map 1 (blue) resembles the N1 component with posterior negativity and fronto-central positivity. Map 2 (green) resembles the P3a component, characterized by fronto-central positivity. Map 3 (red) appears as a hybrid between Map 2 and Map 5. Map 4 (turquoise), defined by bilateral posterior positivity, is clearly identified in the blocked condition but not in the random condition. Map 5 (purple) resembles the P3b component with centroparietal positivity.

**Figure 4.**
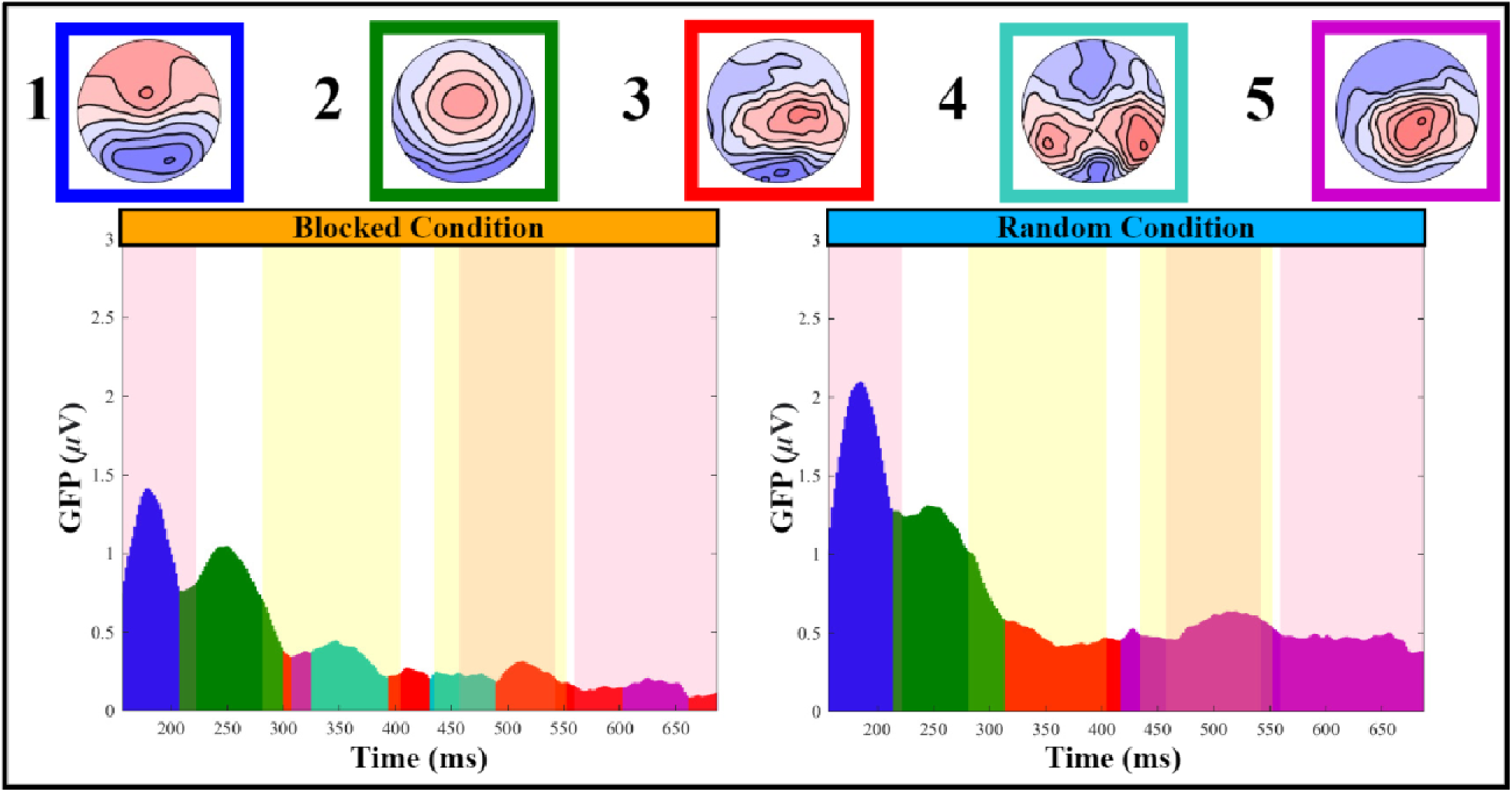
Microstates analysis. The four numbered microstate maps are represented as follows: Map 1 (blue), Map 2 (green), Map 3 (red), Map 4 (turquoise), and Map 5 (purple). The figure includes two graphs showing GFP in microvolts over time (milliseconds) for blocked (left) and random (right) conditions. GFP curves are color-coded to match the microstate maps. Yellow-shaded areas highlight significant time periods (p<0.01) identified by TANOVA analysis. Pink shaded areas highlight significant time periods (p<0.01) identified by GFP analysis.

Map 1 is present in both conditions but shows higher GFP in the random condition, whereas Map 2 is also present in both conditions but shows no significant GFP differences between them. Map 3, absent in the blocked condition, can be a transitional topography between Map 2 and Map 5, as some participants in the random condition may still exhibit P3a-like topography (Map 2) while others may have already transitioned to P3b-like topography (Map 5). This transition may also be present in the blocked condition but, after Map 2, the analysis failed to assign a stable Map in this condition because of the low GFP. Finally, Map 5 appeared consistently only in the random condition, showing both topographical and GFP differences during this period.

The detailed dynamic representation of the topographies supports previous analyses (Figure 5). From 156 ms to 223 ms, both conditions showed similar activity patterns, with a stronger N1-like activi y in the random condition. From 223 to 289 ms, during the P3a-like period, no differences were observed between the groups. From 289 to 422 ms, a transition from a P3a-like to a P3b-like period was evident in the random but not in the blocked condition. Finally, until the end, participants in the random condition displayed strong P3b-like posterior positivity, which was not evident in the blocked condition.

**Figure 5.**
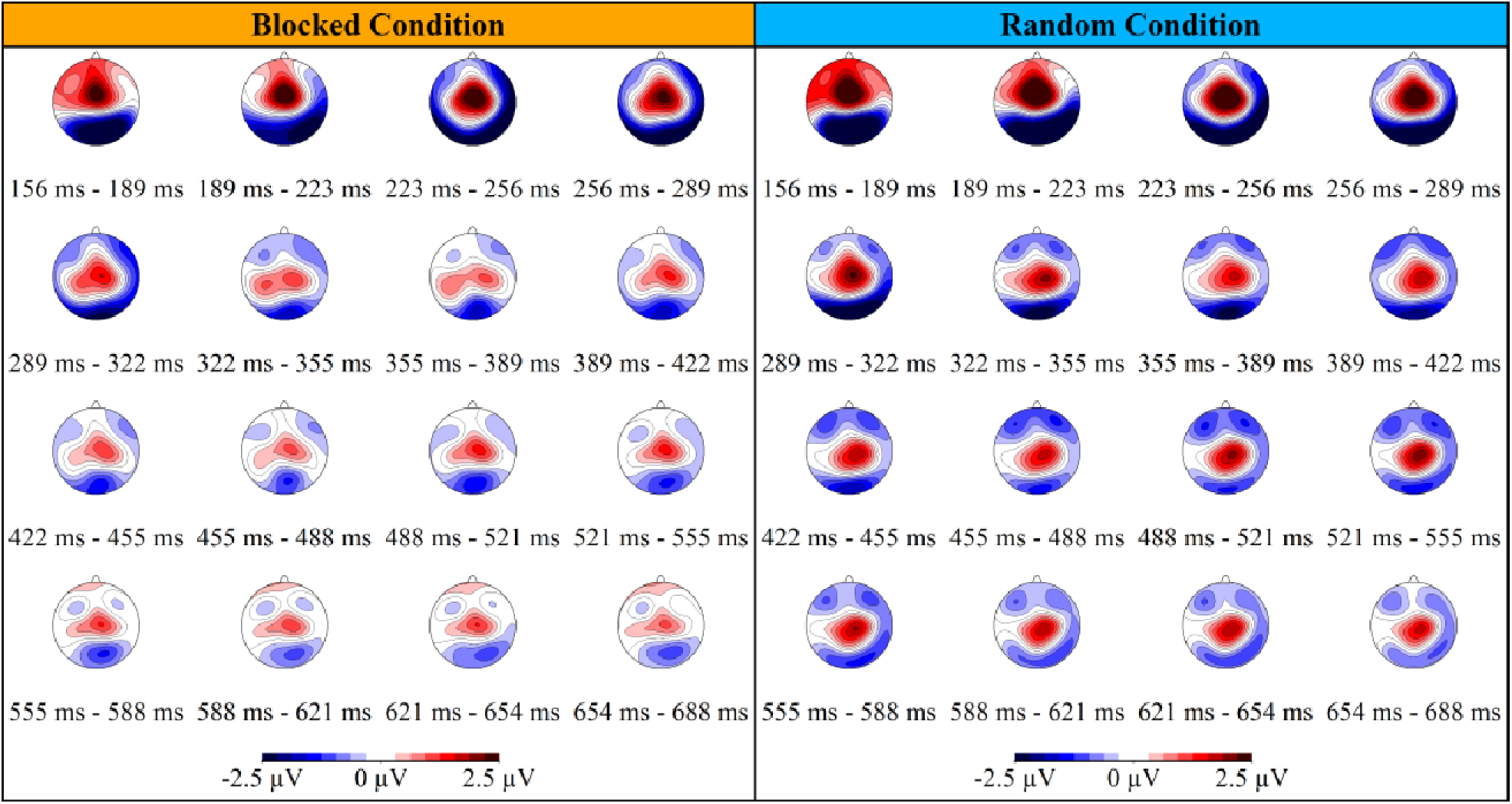
Time-course description of the topographies for blocked (left) and random (right) conditions. The color-coded map illustrates the electrical scalp field potentials, in microvolts, with red indicating positive electrical potential areas and blue indicating negative electrical potential areas.

We then computed standard t-maps to contrast the average topographies of the random and blocked conditions. The t-map for the interval from 156 to 222 ms (Figure 6A) revealed a greater N1-like negative deflection in the random condition than in the blocked condition over the occipital and parietal scalp sites. Consistently, a larger deflection under random conditions was observed at location Pz (Figure 7C, blue curve). During the 281–404 ms interval (Figure 6B), P3a-like frontocentral positivity and posterior negativity were enhanced in the random condition. Higher microvolt amplitudes at the Fpz electrode at the beginning of the period suggest a longer P3a-like topography duration, whereas increased amplitudes at the Pz electrode at the end of the period (Figure 7B) indicate the beginning of the P3b-like component in the random condition. This finding supports the idea that this period marks a transition between processes indexed by P3a and P3b under random conditions. Finally, there was increased P3b-like posterior positivity in the random condition compared to the blocked condition during both the 434–552 ms (Figure 6C) and 559–687 ms periods (Figure 6D). The P3b component was also evident in the amplitude of the Pz electrode under random conditions but not in the blocked condition (Figure 7C, blue curve). The display of all electrodes during the entire epoch for both conditions is available in Supplementary Material (Figure S2).

**Figure 6.**
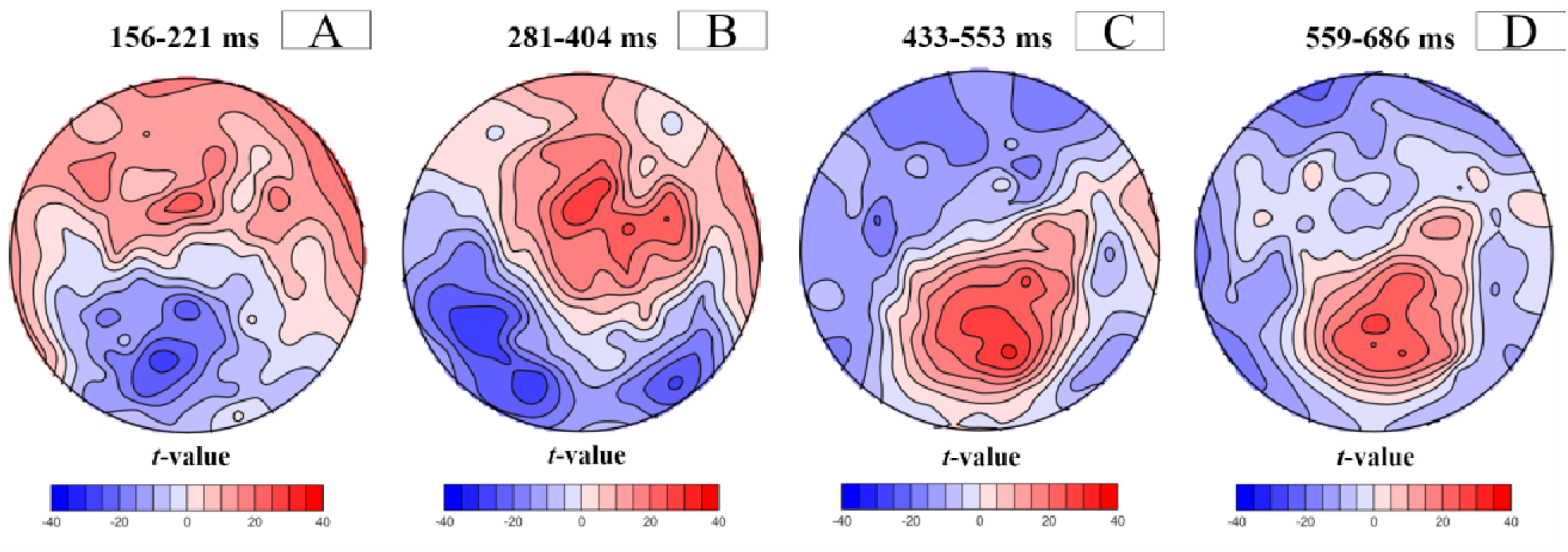
Standard *t*-maps contrasting random and blocked topographies at 156-221 (A), 281-404 (B), 433-553 (C), 559-686 (D) ms periods. Positive (red) and negative (blue) t-values indicate more positive and negative potentials in the random condition than in the blocked condition, respectively.

**Figure 7.**
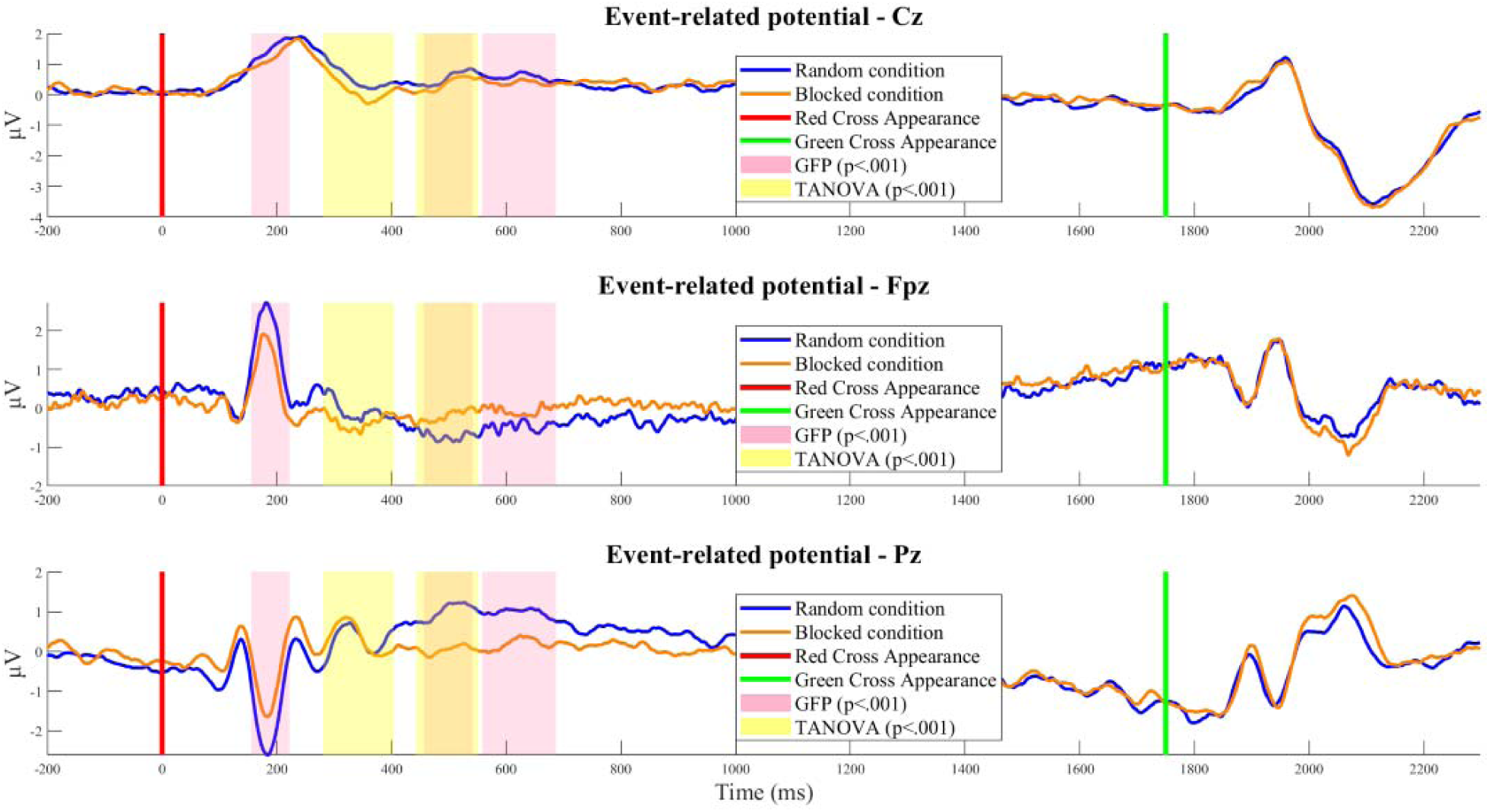
Exemplar ERPs at the Cz, Fpz, and Pz electrodes for the random (blue) and blocked (orange) conditions. The red line indicates when the distance is displayed, and the green line marks the imperative stimulus signaling participants to aim at the target. Yellow-shaded areas indicate periods where TANOVA’s p-value falls below p<.001 and survives post-hoc duration tests. Rose-shaded areas indicate periods where GFP’s p-value falls below p<.001and surives post-hoc duration tests.

### 3.3 Source visualization

The general pattern of activity (Figure 8) was similar between the two conditions until 238 ms, with a succession of centroposterior N1-to a frontocentral P3a-like patterns of activity differing only in the strength of activity. After 238 ms, in the random condition, frontocentral P3a-like activity remained more visible and transitioned to a centroposterior P3b-like pattern of activity that persisted until the end. In contrast, in the blocked condition, a weaker and indistinct pattern of activity was visible from 238 ms until 523 ms, when central activity began and continued until the end.

**Figure 8.**
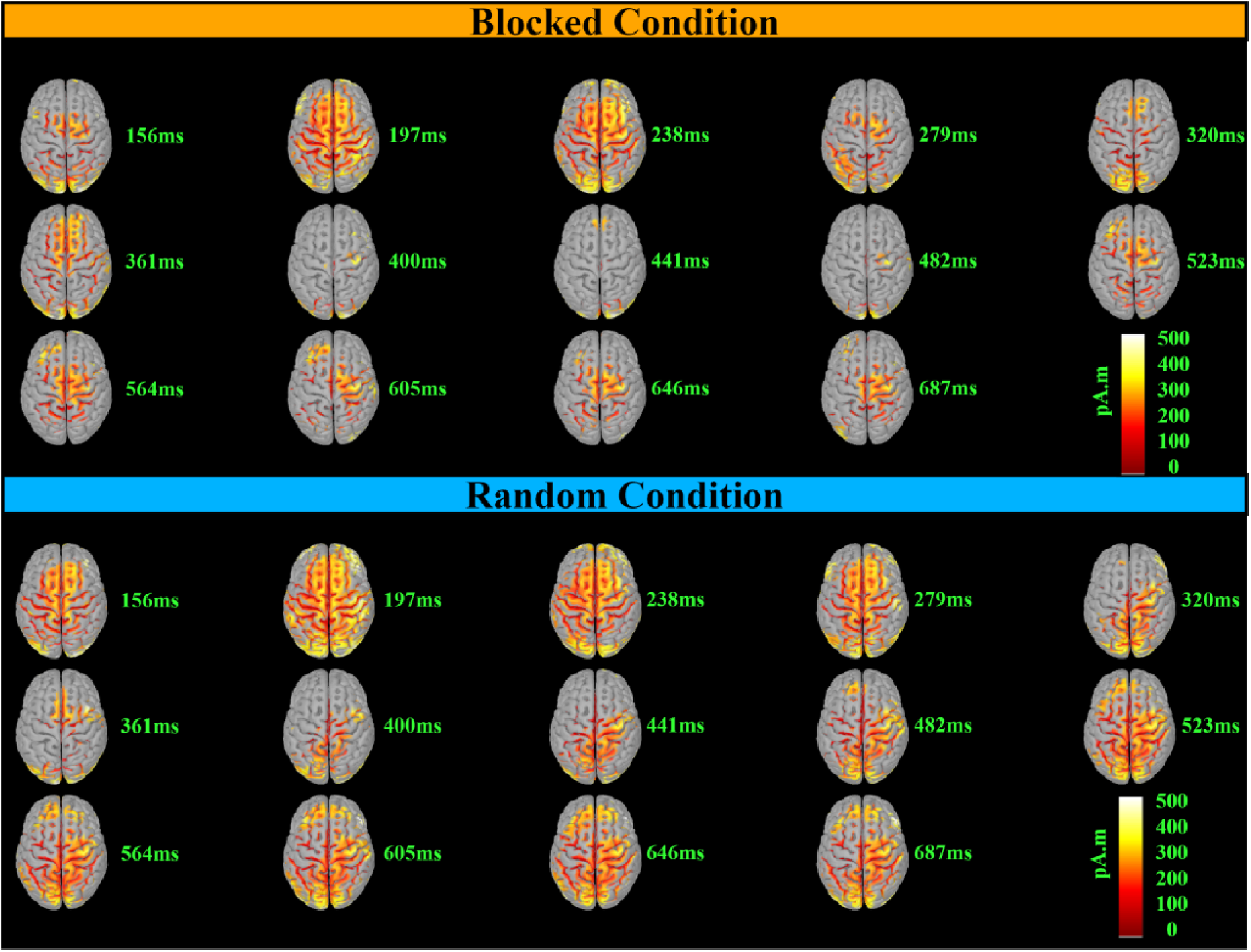
Detailed time-course of the dynamic source estimations for the blocked (top) and random (bottom) conditions. These estimations were computed using a t-test against zero at p<0.01, with FDR correction. The color-coded map illustrates the direction and magnitude of neural activity different from zero, measured in picoamperes times meters (pA·m). Note that this representation does not indicate the directionality of the activity (i.e., whether the potential is positive or negative).

Regarding the neural origins of these differences between the two conditions, the 156–222 ms window (Figure 9) reveals two distinct patterns of increased activity in the random condition compared to the blocked condition. The first pattern demonstrated enhanced activity in the dorsal brain regions, specifically the superior parietal lobe, intraparietal sulcus, precuneus, central cortical region, and superior frontal cortex. The second pattern showed increased ventral activity, encompassing regions such as the lingual gyrus; fusiform gyrus; inferior, middle, and superior temporal gyri; temporal pole; entorhinal cortex; parahippocampal gyrus; pars opercularis; medial and lateral orbitofrontal cortex; insula; and right pars orbitalis.

**Figure 9.**
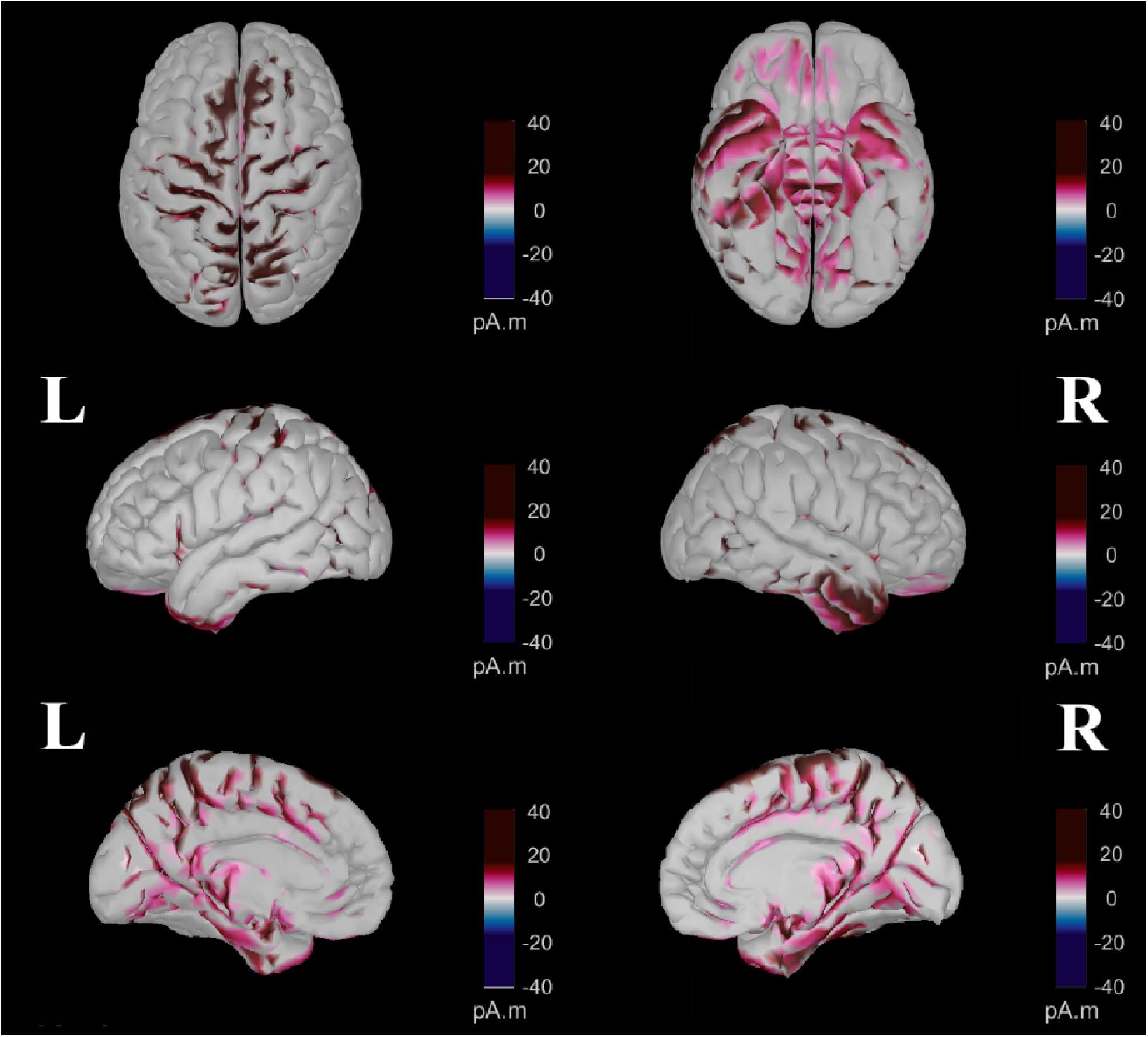
Contrast between random and blocked conditions within the 156-221 ms period computed after parametric statistics (t-test, p<0.01, with FDR correction). The color-coded map indicates the direction and magnitude of differences in neural activity, measured in picoamperes times meters (pA.m). Purple represents areas where there was a significantly more positive electrical potential (higher pA.m values) in the random condition than in the blocked condition. Conversely, blue denotes areas showing significantly more negative electrical potentials in the random condition relative to the blocked condition.

As shown in Figure 10, for the 281–404 ms interval, three distinct patterns of increased activity were observed in the random condition compared to the blocked condition. These patterns included enhanced activity in the bilateral superior midline portions of the precentral, paracentral, and postcentral gyri; elevated activity in the caudal anterior, posterior, and isthmus cingulate cortices; and increased activity in the right inferior and medial temporal gyri, right entorhinal, and right parahippocampal cortices. Conversely, the blocked condition demonstrated higher activity in the right lateral occipital region and bilateral superior frontal cortices.

**Figure 10.**
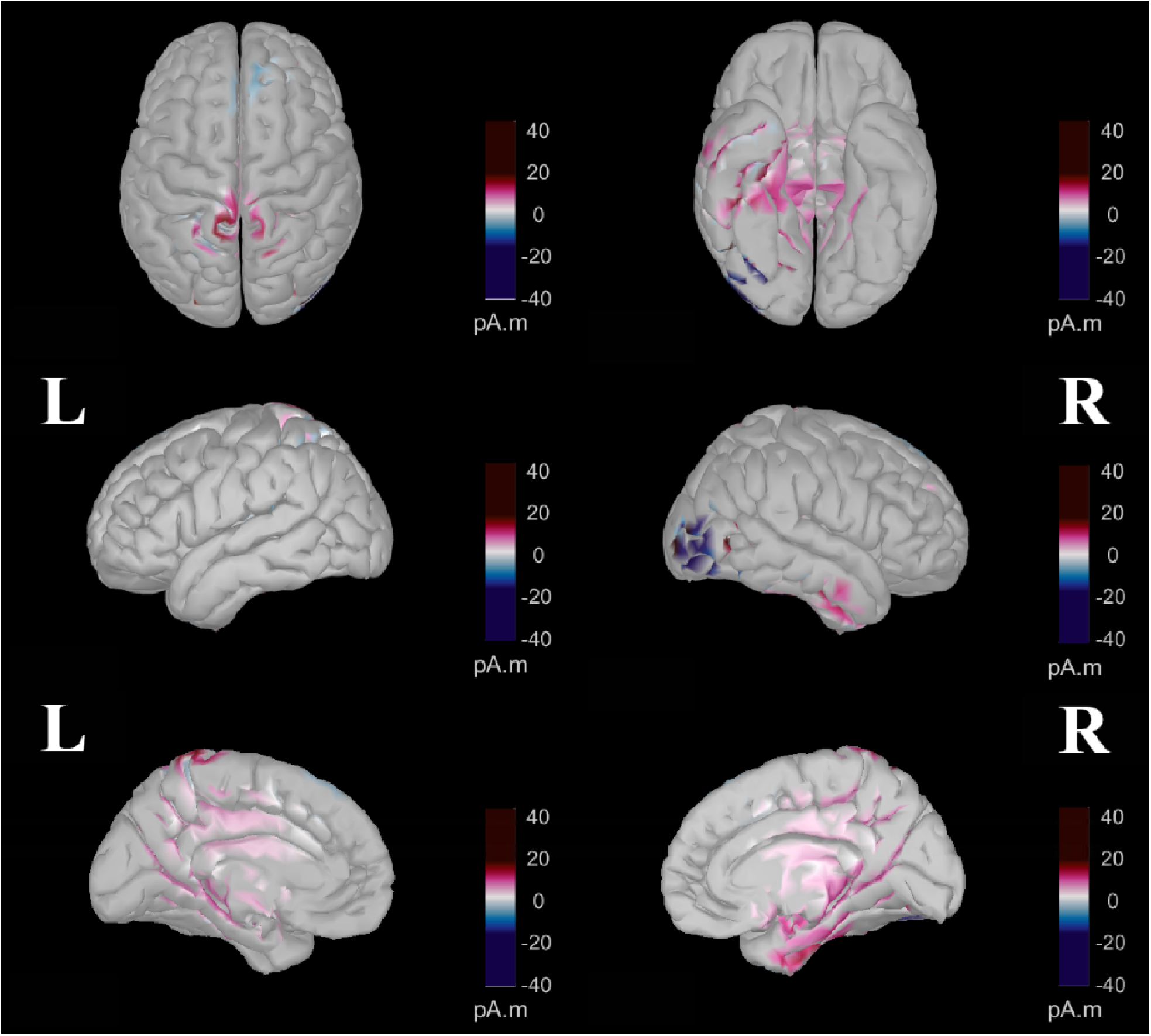
Contrast between random and blocked conditions within the 281–404 ms period computed after parametric statistics (t-test, p<0.01, with FDR correction). The color-coded map indicates the direction and magnitude of differences in neural activity, measured in picoamperes times meters (pA.m). Purple represents areas where there was a significantly more positive electrical potential (higher pA.m values) in the random condition than in the blocked condition. Conversely, blue denotes areas showing significantly more negative electrical potentials in the random condition relative to the blocked condition.

In Figure 11, two distinct patterns of increased activity in the random condition are observed during the 434–552 ms interval. The first pattern is characterized by bilateral activity in the superior parietal cortex, intraparietal junction, and anterior part of the inferior parietal cortex as well as increased activity in the lingual gyrus, precentral gyrus, paracentral lobule, postcentral gyrus, and precuneus. The second pattern involves the activity of the caudal anterior, posterior, and isthmus regions of the cingulate cortex. Additionally, this period showed widespread enhancements in the random condition in both frontal and temporal regions. Conversely, the blocked condition exhibited higher activity in the bilateral cuneus and the right lateral occipital cortex compared to the random condition.

**Figure 11.**
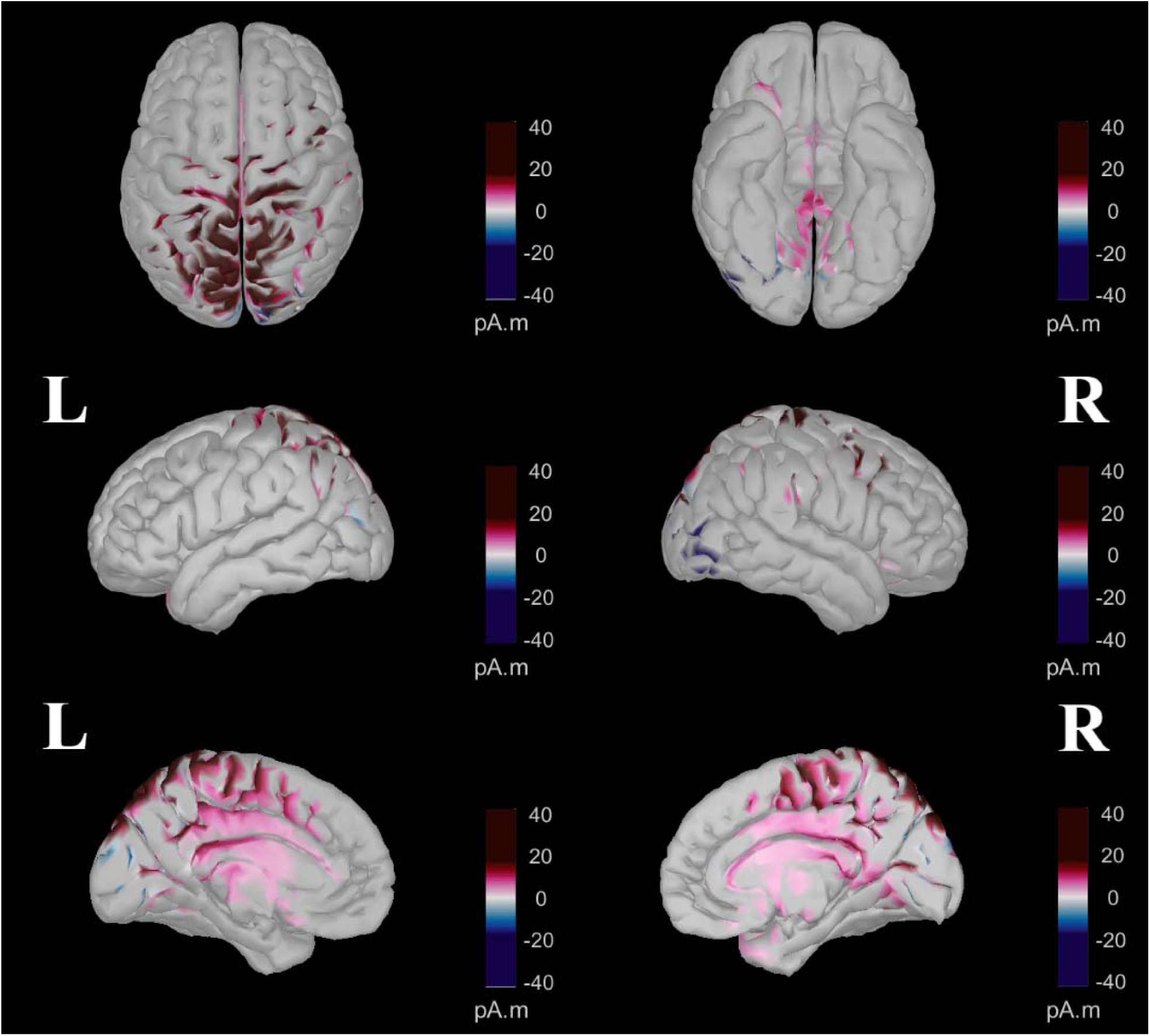
Contrast between random and blocked conditions within the 434–552 ms period computed after parametric statistics (t-test, p<0.01, with FDR correction). The color-coded map indicates the direction and magnitude of differences in neural activity, measured in picoamperes times meters (pA.m). Purple represents areas where there was a significantly more positive electrical potential (higher pA.m values) in the random condition than in the blocked condition. Conversely, blue denotes areas showing significantly more negative electrical potentials in the random condition relative to the blocked condition.

### 3.4 Connectivity analyses

After conducting NBS analyses with paired unidirectional t-tests for each frequency band (theta: whole epoch, alpha: -200–1100 ms, beta and gamma: 0–800 ms) and for each direction, two networks emerged (p < 0.01). These networks are significantly more synchronized in the random than in the blocked condition: one in the theta band (p = 0.006), including 20 edges linking 16 nodes, and the other in the alpha band (p = 0.006), including 18 edges linking 15 nodes. Graphical representations are shown in Figures 12 and 13 along with the regions of the network and the t-statistics between these regions in Tables 1 and 2, respectively.

**Figure 12.**
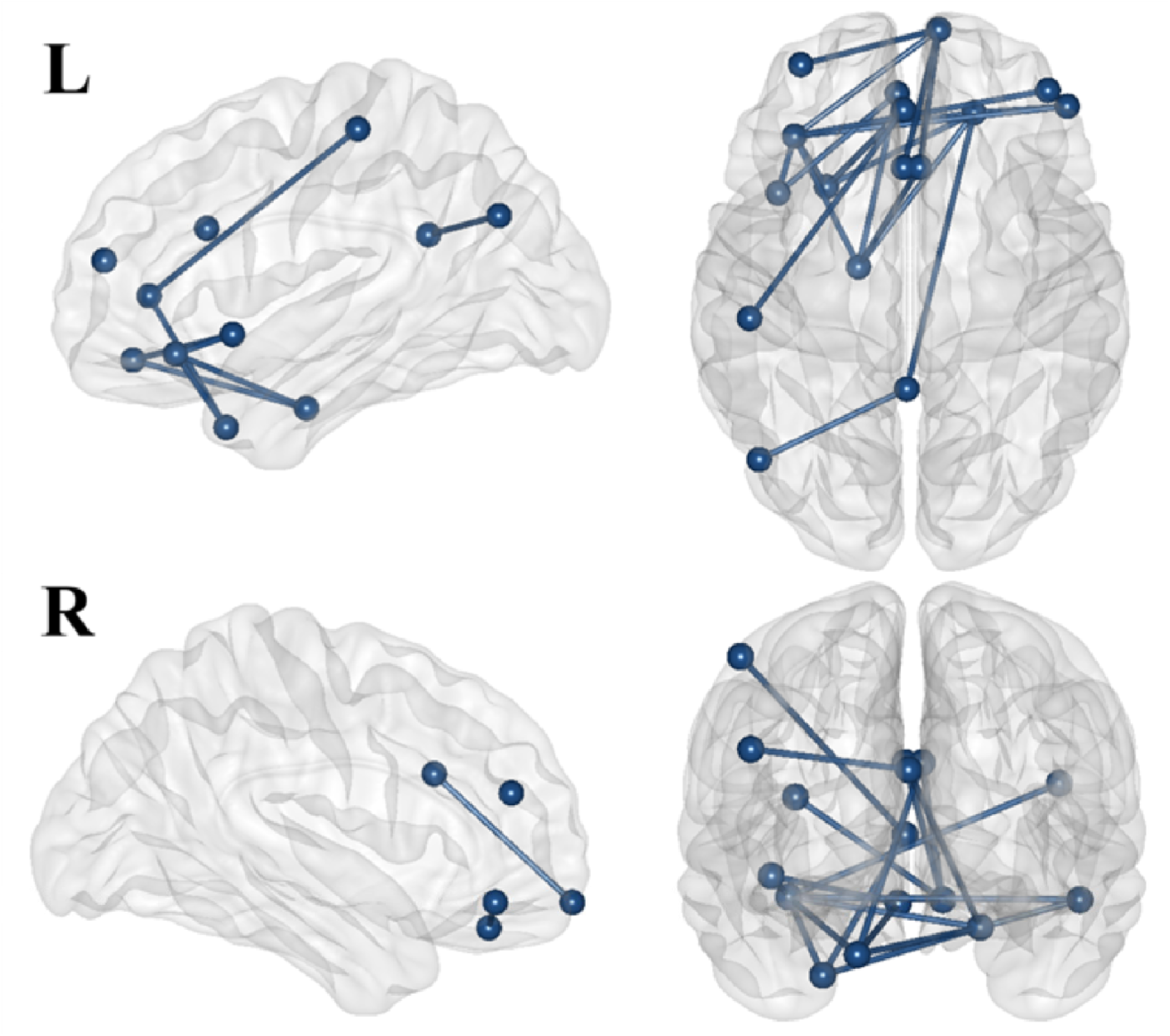
Theta Band Network Connectivity: Random vs. blocked conditions. This figure shows the significant network of nodes linked by edges identified with the unidirectional paired t-test (random > blocked) within the theta frequency band over the whole epoch (period from -200 ms to 2300 ms). R and L refer to the right and left sides, respectively.

**Figure 13.**
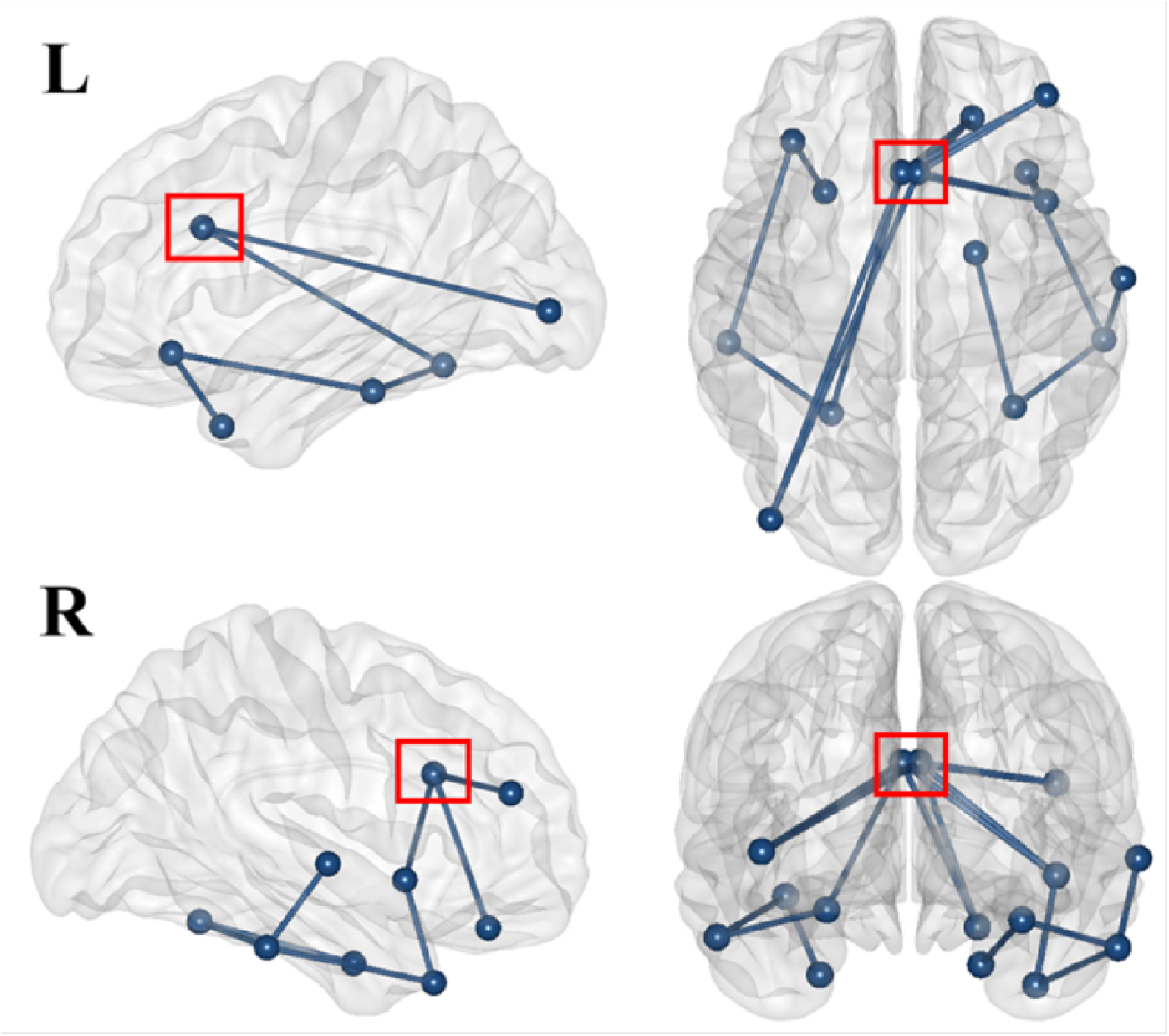
Alpha Band Network Connectivity: Random vs. blocked conditions. This figure shows the significant network of nodes linked by edges identified with the unidirectional paired t-test (random > blocked) within the theta frequency band over the whole epoch (period from -20 ms to 1100 ms). R and L refer to the right and left sides, respectively. The red square is around the anterior cingulate cortex that is the central hub of both our networks.

**Table 1.** Inter-Regional Theta Band Associations. This table lists pairs of regions (Two first columns from the right) along with the corresponding t-statistics (third column), quantifying their association strength in the theta frequency band during the epoch period (-200 ms to 2300 ms). The t-statistics provide a measure of the relationship strength without indicating the directionality of the association.

| Region A | Region B | t-value |
| --- | --- | --- |
| Caudal anteriorcingulate_R | Entorhinal_L | 3.25 |
| Caudal anteriorcingulate_L | Frontalpole_R | 4.03 |
| Caudal anteriorcingulate_R | Frontal pole_R | 3.95 |
| Inferior parietal_L | Isthmus cingulate_L | 3.32 |
| Entorhinal_L | Lateral orbitofrontal_L | 3.47 |
| Frontal pole_R | Lateral orbitofrontal_L | 3.3 |
| Insula_L | Lateral orbitofrontal_L | 3.25 |
| Entorhinal_L | Lateral orbitofrontal_R | 4.32 |
| Isthmus cingulate_L | Lateral orbitofrontal_R | 3.2 |
| Lateral orbitofrontal_L | Lateral orbitofrontal_R | 3.54 |
| Entorhinal_L | Medial orbitofrontal_L | 3.8 |
| Insula_L | Medial orbitofrontal_L | 3.73 |
| Lateral orbitofrontal_L | Pars orbitalis_R | 4.46 |
| Lateral orbitofrontal_R | Pars orbitalis_R | 3.4 |
| Postcentral_L | Rostral anteriorcingulate_L | 3.2 |
| Frontal pole_R | Rostral middlefrontal_L | 3.31 |
| Lateral orbitofrontal_L | Rostral middlefrontal_R | 3.72 |
| Lateral orbitofrontal_L | Temporal pole_L | 3.81 |
| Lateral orbitofrontal_R | Temporal pole_L | 3.29 |
| Rostral anterior cingulate_L | Temporal pole_L | 3.36 |

Theta network analysis (Figure 12; Table 1) revealed a structured connectivity pattern primarily within the prefrontal cortex and extending into the cingulate cortex, with further connections to the temporal and parietal regions. Within the prefrontal cortex, the lateral orbitofrontal cortex emerged as a central node, exhibiting bilateral connections with the frontal pole and lateral orbitofrontal cortex on both sides. This region also showed links to the medial orbitofrontal cortex and pars orbitalis on the right side along with connectivity to the rostral middle frontal region across both hemispheres. Moreover, the right frontal pole has additional connections to the left lateral orbitofrontal cortex. Additionally, the left insula is linked to the prefrontal regions, particularly to the lateral and medial orbitofrontal cortex on the left side. The cingulate cortex was identified as another hub with connections from the anterior cingulate cortex to the left temporal pole and from the left isthmus of the cingulate cortex to the left inferior parietal lobule, highlighting the integration of the cingulate cortex as a hub between the frontal and parietal regions. Temporal lobe connectivity included the left entorhinal cortex, which connects the right lateral orbitofrontal cortex and the left medial orbitofrontal cortex.

The alpha network (Figure 13; Table 2) was centered around the anterior cingulate cortex, linking the ventral and prefrontal regions. The ventral areas included the left lateral occipital cortex, bilateral fusiform gyrus, bilateral inferior and right superior temporal regions, bilateral temporal pole, and the right entorhinal cortex. Prefrontal connections involved the bilateral insula, the right lateral orbitofrontal cortex, and the right rostral middle frontal region.

**Table 2.** Inter-Regional Alpha Band Associations. This table lists pairs of regions (Two first columns from the right) along with the corresponding t-statistics (third column), quantifying their association strength in the theta frequency band during the epoch period (-200 ms to 1100 ms). The t-statistics provide a measure of the relationship strength without indicating the directionality of the association.

| Region A | Region B | t-value |
| --- | --- | --- |
| Caudal anterior cingulate_L | Caudal anterior cingulate_R | 3.46 |
| Caudal anterior cingulate_L | Fusiform_L | 3.35 |
| Entorhinal_R | Fusiform_R | 3.15 |
| Fusiform_L | Inferior temporal_L | 5.72 |
| Fusiform_R | Inferior temporal_R | 3.77 |
| Caudal anterior cingulate_L | Insula_R | 3.5 |
| Caudal anterior cingulate_R | Insula_R | 3.42 |
| Caudal anterior cingulate_L | Lateral occipital_L | 4.45 |
| Caudal anterior cingulate_R | Lateral occipital_L | 3.82 |
| Inferior temporal_L | Lateral orbitofrontal_L | 3.22 |
| Caudal anterior cingulate_L | Lateral orbitofrontal_R | 3.61 |
| Caudal anterior cingulate_R | Lateral orbitofrontal_R | 4.59 |
| Caudal anterior cingulate_L | Rostral middle frontal_R | 3.91 |
| Caudal anterior cingulate_R | Rostral middle frontal_R | 3.78 |
| Inferior temporal_R | Superior temporal_R | 3.14 |
| Lateral orbitofrontal_L | Temporal pole_L | 3.38 |
| Inferior temporal_R | Temporal pole_R | 3.35 |
| Insula_R | Temporal pole_R | 3.26 |

## 4. Discussion

The study aimed to examine how CI affects the spatial and temporal neural correlates of attention and working memory. Using a high-density EEG multi-scale analysis approach, we investigated topographical patterns, source estimation, and source connectivity. We hypothesized that ERP markers would distinguish random from blocked conditions, with higher N1 and P3a components indicating increased perceptual/attentional demand and larger P3b component indicating greater working memory engagement. We also expected source estimation to reveal brain activity related to both attention and working memory, and the random condition showing enhanced synchronization within the frontoparietal Executive Control Network (ECN).

Our analyses supported these hypotheses. Topographical analyses showed significant differences between conditions from 156 to 687 ms, indexed by variations in the N1, P3a, and P3b components. During the 156-221 ms window, both conditions exhibited a posterior negativity and a negative deflection at the Pz electrode characteristic of N1, but with greater amplitude in the random condition. From 220 to 280 ms, both conditions showed similar positive activity across frontocentral regions characteristic of P3a (Halgren et al. 1998; Kok 2001; Polich 2003; Huang et al. 2015). However, from 281-404 ms, significant topographical differences occurred. The random condition exhibited a hybrid P3a/P3b-like topography, with successive topographies and ERPs at Fpz and Pz electrodes indicating a transition from P3a-like frontocentral to P3b-like centroparietal activity; a transition not observed in the blocked condition. This suggests that the P3a component lasts longer in the random condition and transitions to P3b. Indeed, from 434 to 687 ms, only the random condition showed strong parietal positivity and a deflection at Pz electrode characteristic of the P3b component (Polich 2007; Verleger 2020). In summary, topographical analyses suggest that both conditions involve the same neural generators until 280 ms, with higher N1 activity in the random condition. Afterwards, different neural generators are involved, with a longer P3a in the random condition and notable reduction of an identifiable P3b in the blocked condition.

The source visualization confirmed the general pattern observed with topographies. Until 238 ms, the pattern of activity was similar between conditions but stronger in the random condition. After 238 ms, activity in the random condition transitioned from frontocentral to centroposterior regions, while the blocked condition showed weaker activity. An analysis of the average activity contrast between conditions from 156-221 ms confirmed significantly greater activity in the random condition, particularly in the posterior regions and the ventral pathway. This includes the lingual and fusiform gyri, temporal gyri, entorhinal cortex, medial and lateral orbitofrontal cortex, and insula. This pattern aligns with the N1 component, associated with perceptual processes, and believed to originate in the occipital and posterior parietal cortices and the ventral stream (Murray et al. 2002; Murray et al. 2004; Sehatpour et al. 2006). From 281-404 ms, the random condition exhibited greater activity in the parietal cortex and the anterior and posterior cingulate cortex, regions known to be the neural generators of the P3a component and associated to attentional processes (Halgren et al. 1998; Crottaz-Herbette and Menon 2006; Volpe et al. 2007; Huang et al. 2015). From 434-552 ms, the random condition triggered significantly more activity in the parietal and cingulate cortices, including the caudal anterior, posterior, and isthmus regions, linked to the P3b component and working memory (Polich 2003; Bledowski et al. 2004; Polich and Criado 2006; Polich 2007; Volpe et al. 2007; Menon and Uddin 2010). To summarize, source analyses confirmed reduced activity after P3a in the blocked condition, with strong posterior activity in the random condition. Differences at N1 suggest greater ventral regions activity in the random condition, while later differences between conditions involve more cingulate and parietal cortex activity in the random condition.

Finally, the source connectivity analysis confirmed greater theta synchronization within the frontoparietal ECN in the random condition. The theta network, observed throughout the entire epoch, exhibited a structured connectivity pattern primarily involving the prefrontal cortex and extending into the cingulate cortex, with additional links to temporal and parietal regions. Additionally, an alpha network with a ventral connectivity pattern was observed from -200 to 1100 ms, centered around the anterior cingulate cortex and connected to various prefrontal regions, including the lateral occipital cortex, fusiform gyrus, temporal regions, insula, and orbitofrontal cortex.

### Modulation of Attention and Perception-Related Processes by CI

The visual N1 component, present in both conditions but more pronounced in the random condition, indicates increased stimulus processing and visuospatial attention in the random condition (Correa et al. 2006; Krigolson et al. 2015). The N1 component is observed in the hemisphere contralateral to the attended stimulus (Harter et al. 1989; Kasai et al. 2003) and facilitates attention orientation and processing of task-relevant stimuli (Luck et al. 1990; Slagter et al. 2016). It is associated with the discrimination process, as shown by Vogel and Luck (2000) and Hopf et al. (2002) who found higher N1 amplitude in tasks requiring discrimination between colors or shapes (letters) compared to simple reaction tasks. N1 amplitude also increases with the difficulty of discrimination, such as when non-targets resemble targets (Fedota et al. 2012; Roberts et al. 2014). This could explain the higher N1 observed in the random condition of our study, where participant had to discriminate between distances in each trial, unlike in the blocked condition, where they passively viewed the same distance requiring no discrimination.

During the same period, we observed greater activity in the random than in the blocked condition in ventral regions. Interestingly, the visual N1 component is described to originate within the ventral stream (Doniger et al. 2000; Vogel and Luck 2000; Doniger et al. 2002; Hopf et al., 2002; Slagter et al. 2016; Vogel & Luck, 2000), which, along with the dorsal stream, constitutes one of the two main visual processing pathways. These pathways are often referred to as the “what” and “where” streams, respectively (Milner and Goodale 2008; David Milner 2015). The occipitotemporal ventral stream processes visual information for object perception, categorization, and recognition, using invariant features such as shape, color, texture, and spatial relationships. Conversely, the occipitoparietal dorsal stream focuses on transforming visual information into action coordinates, guiding movements and spatial localization for the visual control of skilled actions like reaching and grasping, without directly contributing to object categorization (Milner & Goodale, 2008; Milner 2015). Given the nature of our task, which involves aiming at a target based on its spatial relationship with different starting positions in 2D space, one might initially expect to see a condition differences in the involvement of the dorsal stream. However, while the dorsal stream can encode spatial relationships egocentrically (centered around the viewer for immediate actions), the ventral stream can process them allocentrically (using an external frame of reference) (Schenk 2006; Zaehle et al. 2007; Sereno and Lehky 2011). The entorhinal cortex, more active in the random compared to the blocked condition, plays a critical role in spatial navigation and memory. It contains grid cells that encode spatial information in allocentric terms, offering a two-dimensional map-like representation of the environment (Fyhn et al. 2004; Hafting et al. 2005; Constantinidis and Klingberg 2016; Nau et al. 2018; Nadasdy et al. 2022). Additionally, the coding of distances extends beyond exact metric to categorical terms like above/below or right/left comparisons (Kosslyn et al. 1992; Noordzij and Postma 2005; Baumann et al. 2012). Tasks that are two-dimensional and focus on a small region of space, like our aiming task, tend to promote categorical over coordinate metric judgment (Borst and Kosslyn 2010; Ruotolo et al. 2016). Thus, the random condition, with its changing distances, could require greater perceptual discrimination and categorization within the ventral stream.

The anterior cingulate cortex (ACC) emerged as the principal hub in the identified alpha network, during the –200 to 1100 ms period, and source estimation analysis within the 281-404 ms window. This region is also believed to be a significant generator of the P3a component (Halgren et al. 1998; Polich 2003; Crottaz-Herbette and Menon 2006; Polich and Criado 2006; Polich 2007; Volpe et al. 2007; Huang et al. 2015). The P3a component is primarily associated with the involuntary attentional shift towards novel, unexpected, or salient stimuli, indexing the brain’s modulation of the allocation of cognitive resources to assess their significance and to enhance attentional allocations (Escera et al. 2000; Friedman et al. 2001; Polich & Comerchero 2003; Polich 2007). Moreover, P3a reflects shifts in attention triggered by both task-relevant and task-irrelevant changes in the stimulus attributes (Squires et al. 1975; Barceló et al. 2002; Berti 2008; Hölig and Berti 2010; Berti 2016). For example, in a Wisconsin Card Sorting Test (WCST), where subjects were asked to sort cards according to varying criteria (color, shape, or number) and adapt to changes in these sorting rules, it has been shown that switching from one sorting criterion to another elicits a greater frontal P3a response (Barceló et al. 2002). This is congruent with our study, where we observed higher P3a amplitudes in the random condition, requiring participants to switch the aiming distance in each trial, compared to the blocked condition, in which the aiming distance remained constant. Furthermore, this interpretation is reinforced by the increased activity observed in the random condition within the same period in the ACC and the right DLPFC. These regions have been linked to task switching, with the ACC playing a crucial role in monitoring attentional shifts during task switching, and the DLPFC essential for maintaining cognitive goals and managing distractions from competing tasks (Luks et al. 2002; Kondo et al. 2004; Liston et al. 2006; Hyafil et al. 2009).

Our hypothesis of higher perceptual/attentional processing during the random condition is supported by the large-scale networks more synchronized in the random condition compared to the blocked condition in the alpha band. Alpha oscillations are linked to attentional functions, such as enhancing task-related sensory processing by inhibiting irrelevant information (Foxe and Snyder 2011; Klimesch 2012; Van Diepen et al. 2019), and maintaining tonic alertness (Sadaghiani et al. 2010; Huang et al. 2023). This ability to maintain tonic alertness has been attributed to the Cingulo-Opercular Network, including the anterior cingulate cortex, operculum, and insula (Sadaghiani and D’Esposito 2015; Coste and Kleinschmidt 2016). The involvement of the anterior cingulate cortex, a hub in our identified network, and the Insula suggests that the random condition required consistently higher alertness than the blocked condition. This heightened alertness would improve detection and response to changing task requirements, unlike the blocked condition, where such detection is less needed.

### Modulation of memory working-related processes by CI

P3a is also involved in working memory processes. In a study by Berti (2016), participants engaged in a memory-updating task with three phases: presentation, updating, and recall. They memorized four digits, then either replaced a digit or adjusted it based on cues. Switching attention to a new digit resulted in longer updating times and a higher P3a amplitude, indicating P3a’s role in reallocating attention within working memory (Berti 2008; Berti 2016). After detecting a distracting stimulus, as reflected by the P3a, task-relevant information processing follows, indexed by the P3b component (Polich 2003; Crottaz-Herbette & Menon 2006; Polich 2007).

P3b is typically observed in three-stimulus oddball tasks, where participants respond to infrequent target stimuli among regular non-targets. P3b amplitude is higher for infrequent targets, indicating task-relevant information processing. Two main hypotheses explain P3b: the working memory update hypothesis (Polich 2003; Polich & Criado 2006; Polich 2007) and the stimulus-response link reactivation hypothesis (Verleger 2008; Verleger et al. 2015; Verleger 2020). The working memory update hypothesis suggests P3b component is deeply intertwined with working memory mechanisms. This model suggests that P3b is elicited when the brain updates the context within working memory in response to a mismatch, reflected in P3a, between an expected schema and an actual task-relevant stimulus, and reflects the need to store rare or significant stimuli in memory for future recognition (Polich 2003; Polich & Criado 2006; Polich 2007). In contrast, the stimulus-response link reactivation hypothesis emphasizes P3b’s role in linking perception to action, reflecting the reactivation of stimulus-response links. These links, once established through repetition, require reactivation in tasks where they are not repeated in each trial. The modulation of P3b amplitude in response to less frequent stimuli highlights the greater neural effort involved in reactivating these associations (Verleger 2008; Verleger et al. 2015; Verleger 2020).

Both hypotheses can explain the higher P3b observed in the random condition compared to the blocked condition. According to the working memory update hypothesis, encountering a new distance in the random condition leads the brain to detect a mismatch from the previous trial, reflected by a higher P3a, followed by a P3b, reflecting the update of the currently requested distance schema in working memory and the storage of this information for future use. Alternatively, the stimulus-response link reactivation hypothesis suggests that participants in the random condition need to discard the previous stimulus-response mapping and reactivate the new one for the current trial, reflected in the higher P3b.

Our hypothesis that mnemonic processing is enhanced during the random condition is also supported by our findings of greater synchronization in the theta band network throughout the entire epoch in the random condition compared to the blocked condition. This network, resembling the Executive Central Network (ECN), includes interconnections within the frontal and prefrontal regions, and between these regions and the inferior parietal cortex, ACC, insula, and temporal brain areas (Collette and Van der Linden 2002; Vincent et al. 2008; Niendam et al. 2012). The frontoparietal network is linked to cognitive functions such as attentional control, flexibility, inhibition, and working memory (Niendam et al., 2012) and is associated with enhanced executive function performance (Shen et al. 2019). Working memory training has been shown to induce changes in ECN (Langer et al. 2013; Zhang et al. 2015; Thompson et al. 2016), and frontoparietal activity during tasks differentiates good from poor performers (Nyberg et al. 2009). The ECN’s reliance on theta band connectivity is also well-documented, with increased frontoparietal theta synchronization during working memory tasks correlating with better performance (Mizuhara and Yamaguchi 2007; Gulbinaite et al. 2014; Kawasaki et al. 2014; van Son et al. 2019; Xiong et al. 2024). Finally, this relationship is supported by evidence that transcranial direct current stimulation enhances frontoparietal theta synchronization and improves working memory (Violante et al. 2017).

## Conclusion

Collectively, the results of our multi-scale analyses are compatible with the hypotheses that the random condition induces greater attentional and perceptual demands, as well as increased working memory engagement, compared to the blocked condition. The enhanced topographical N1-like component in the random condition, along with greater activity in the ventral visual pathway, suggests a higher need for stimulus discrimination due to changing distances. Additionally, the extended duration of the P3a-like topography and the greater involvement of the cingulate and prefrontal cortices in the random condition indicate a greater need for attentional and cognitive control to manage the unpredictible shifts in aiming distance. The large-scale connectivity analysis revealed greater alpha synchronization in a network centered around the ACC in the random condition, suggesting consistently higher alertness and attentional engagement beyond the specific periods indexed by the N1 and P3a. In the random condition, we observed a transition from P3a-like to P3b-like topographies, accompanied by a frontocentral to posterior pattern of source activity. This transition may represent a greater reliance on working memory, possibly due to the need to update the distance parameter at each attempt or the reactivation of the stimulus-response link at each trial due to its inconsistent mapping. Furthermore, the persistent large-scale frontoparietal ECN network in the theta band throughout the entire epoch may suggest not only enhanced working memory processes but rather a heightened need for executive function and task-relevant processing across the entire task.

Finally, our findings offer exploratory evidence, leveraging the excellent spatial and temporal resolution of high-density EEG, suggesting that the contextual interference effect is likely not due to a single process unique to the random condition. Rather, it involves multiple processes modulated by persistent large-scale network activity. Further longitudinal studies are needed to investigate how contextual interference affects topographical, sources and network modulation over time.

## ACKNOWLEDGMENTS

We wish to express our gratitude to our colleagues at the Brain Electrophysiology Attention Movement Laboratory for their assistance in the execution of the experiment. A special acknowledgment is due to Elsa Raynal and Francesco Di Muccio for their contributions. Additionally, we are deeply thankful to the participants of the experiment, whose dedication was pivotal to the success of this study.

## CONSENT TO PARTICIPATE

The authors declare that every participant provided written, informed consent prior to participating in the study.

