## Supplementary material for "Enhancing Perceptual, Attentional, and Working Memory Demands through Variable Practice Schedules: Insights from High-Density EEG Multi-Scale Analyses": Table S1

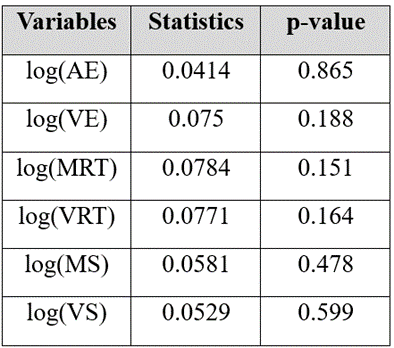

Table S1. Result of the Kolmogorov-Smirnov normality test for each of the 6 log-transformed behavioral variables. P-value above 0.05 indicates the non-violation of the normality hypothesis.
