## Supplementary material for "Enhancing Perceptual, Attentional, and Working Memory Demands through Variable Practice Schedules: Insights from High-Density EEG Multi-Scale Analyses": Figure S2

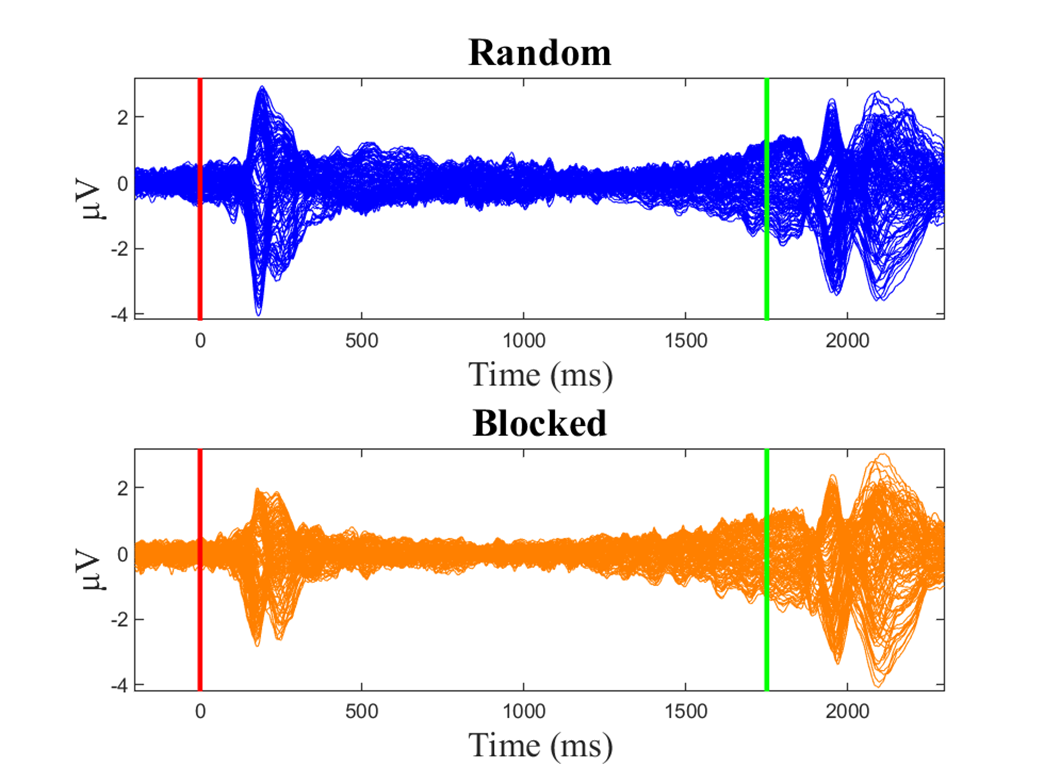

Figure S2. Mean ERPs of all electrodes for random (blue) and blocked (orange) conditions. The red line marks the moment when the distance to be executed is displayed and the green line signifies the imperative stimulus when participants are required to aim at the target.
