## Supplementary material for "Enhancing Perceptual, Attentional, and Working Memory Demands through Variable Practice Schedules: Insights from High-Density EEG Multi-Scale Analyses": Figure S1

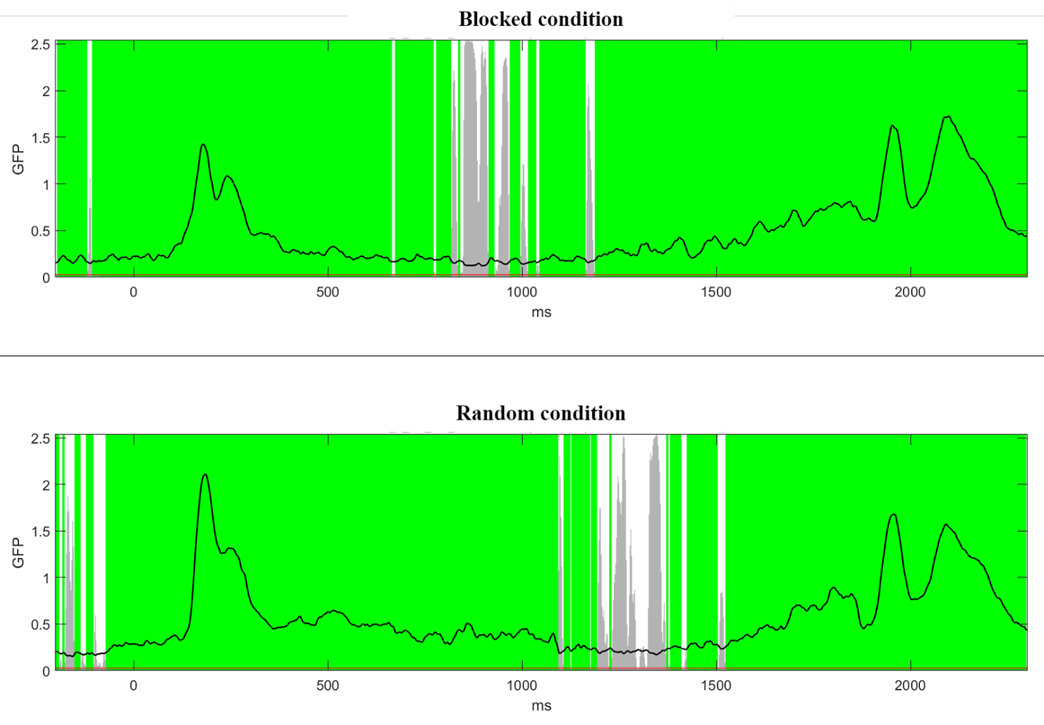

Figure S1. Result of Topographical Consistency Test for each condition. The figure depicts the GFP dynamic over the epoch. The green areas represent period of consistent topographies within each condition. Grey areas represents period of inconsistent topographies within each condition.
